# An A-T Hoogsteen base pair in a naked DNA hairpin motif: A Protein-Recognized Conformation

**DOI:** 10.1101/2024.12.22.630000

**Authors:** Serafima Guseva, Or Szekely, Ainan Geng, Kai Smith, Supriya Pratihar, Stephanie Gu, Hashim M. Al-Hashimi

## Abstract

In duplex DNA, A-T and G-C form Watson-Crick base pairs, and Hoogsteen pairing only dominates upon protein binding or DNA damage. Using NMR, we show that an A-T Hoogsteen base pair previously observed in crystal structures of transposon DNA hairpins bound to TnpA protein forms in solution even in the absence of TnpA. This Hoogsteen base pair, located adjacent to a dinucleotide apical loop, exists in dynamic equilibrium with a minor Watson-Crick conformation (population ∼11% and lifetime ∼55 µs). Extending the apical loop to three residues inverted the equilibrium, making Watson-Crick the dominant state and the Hoogsteen conformation recognized by TnpA a minor state (population ∼14% and lifetime ∼28 µs). The propensity for Hoogsteen pairing depended on apical loop residues, which form contacts directly or indirectly stabilizing the Hoogsteen conformation. A structure survey did not reveal Hoogsteen pairing near RNA apical loops making them unique to DNA. Our results demonstrate that Hoogsteen can be the dominant state even in naked unmodified duplex DNA and identify 5’-CTT(T/C)AG-3’ as a DNA-specific apical loop motif stabilized by Hoogsteen pairing. Hoogsteen base pairs may be prevalent in DNA hairpins forming during replication and transcription, with broad implications for the genomic landscape.

## Introduction

It is well established that the DNA double helix is composed of A-T and G-C Watson-Crick base pairs (bps)^1^. However, it has been known for decades that both A-T and G-C can also pair in an alternative conformation called “Hoogsteen”^2^. Starting from a Watson-Crick bp, the Hoogsteen conformation can be obtained by flipping the purine base 180° from the *anti* to the *syn* conformation and by bringing the two bases into closer proximity by ∼2.5 Å to form a unique set of hydrogen bonds (Figure 1A)^3^. In solution, the bps in naked duplex DNA are predominantly Watson-Crick, and the Hoogsteen conformation only becomes the dominant state when DNA is damaged or bound to proteins or small molecules^4–8^. NMR studies, however, have revealed that even in naked DNA, Watson-Crick bps transiently form the Hoogsteen conformation, though it typically represents less than a few percent of the total population^9–10^. Here, using NMR spectroscopy, we show that an A-T bp adopts the Hoogsteen conformation in solution as the dominant state even in the absence of bound protein or DNA damage.

**Figure 1.**
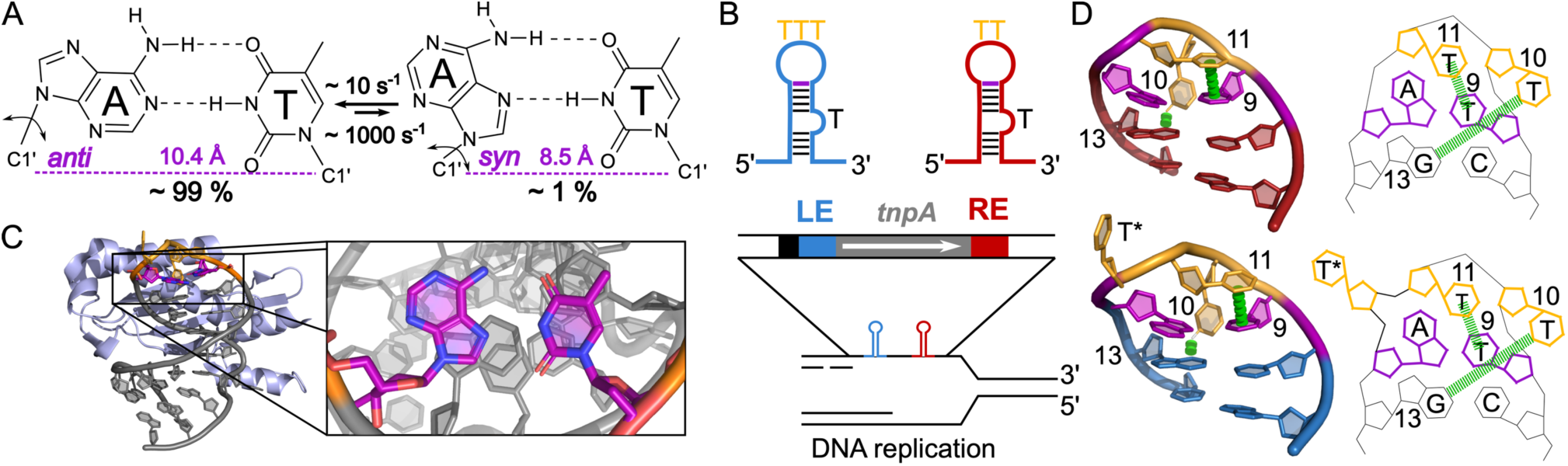
An A-T Hoogsteen base pair is observed robustly adjacent to an apical loop in crystal structures of TnpA transposase in complex with RE and LE hairpins from two members of the IS200/IS605 family. (A) In naked duplex DNA, the A-T Watson-Crick base pair (left) exists in dynamic equilibrium with a short-lived and lowly-populated Hoogsteen conformation (right). (B) Schematic of the IS608 LE and RE hairpins from the IS200/IS605 family containing trinucleotide and dinucleotide thymine apical loops, respectively. The IS608 transposon encodes TnpA transposase and is surrounded by the subterminal hairpins LE (blue) and RE (red), which are formed during replication through folding of single-stranded DNA intermediates. (C) Representative crystal structure (PDB ID: 2VJV) of TnpA IS608 (light blue) in complex with LE hairpin (gray) with an A12-T9 Hoogsteen base pair (purple) forming adjacent to the trinucleotide apical loop. (D) TnpA-bound RE and LE hairpins exhibit an A12(*syn*)-T9 Hoogsteen base pair (purple). Interactions involving apical loop residues (yellow) which may directly or indirectly stabilize the Hoogsteen conformation are highlighted in green.

Our study was inspired by TnpA, a family of bacterial transposase proteins that catalyze the movement of specific DNA sequences, called ‘transposons’, from one genomic location to another. Such transposable elements contribute to genetic diversity and play significant roles in the spread of antibiotic resistance and in the adaptation to new environments^11^. Transposases mediate transposition by specifically recognizing DNA sequences at the ends of the transposon, excising the element from the donor site, and integrating it into a new target site. While many transposases act on double-stranded DNA, TnpA from the insertion sequence family IS200/IS605 targets DNA hairpins formed by single-stranded DNA intermediates during replication (Figure 1B)^12–14^. TnpA specifically recognizes subterminal hairpins (Figure 1B) forming at the left end (LE) and right end (RE) of the insertion sequence^15–17^. In the IS200/IS605 family member IS608, the RE hairpin has a dinucleotide apical loop whereas LE tends to have a trinucleotide apical loop (Figure 1B)^18^.

What was intriguing about TnpA from the IS200/IS605 family is that numerous crystal structures revealed it employed an A-T Hoogsteen bp to bind target DNA hairpins^15–17^. Specifically, there were 11 crystal structures of TnpA-DNA complexes comprising 10 RE and 19 LE hairpins^15–17^ derived from two members of the IS200/IS605 family, IS608 and ISDra2 (Figure S1). In all cases, an A(*syn*)-T Hoogsteen bp was observed adjacent to the dinucleotide (n=19) or trinucleotide (n=10) apical loop (Figure 1C,D). This Hoogsteen bp was observed robustly across varying crystallization conditions and crystal space groups^15–17^. Our own reanalysis of the electron density (see Methods and Figure S2) supports modeling a Hoogsteen conformation at this bp position, helping to rule out modeling errors or ambiguities due to poor electron density reported recently^6, 19^.

What drives this A-T bp to adopt the Hoogsteen conformation when bound to TnpA? Based on the crystal structures, several contacts exist between TnpA and the backbone of the DNA apical loop, but none of them are base-specific contacts that could preferentially stabilize the adenine base in a *syn* versus *anti* orientation. Similarly, no crystal contacts could be identified that stabilize the Hoogsteen conformation. Instead, the A(*syn*)-T Hoogsteen conformation appears to be stabilized by intramolecular DNA interactions within the apical loop. Specifically, T11 stacks directly on the Hoogsteen bp while T10 folds back to form a hydrogen bond with G13(N3) within the duplex, a conformation that might be more easily accommodated by the constricted A(*syn*)-T Hoogsteen bp relative to a Watson-Crick bp (Figure 1D). Thus, we hypothesized that the A(*syn*)-T Hoogsteen bp observed in LE and RE in complex with TnpA transposase might form even in the absence of the bound protein. Such a result would have important implications for the prevalence of Hoogsteen bps in DNA hairpins genome-wide.

## Results

### The A-T Base Pair in RE Predominantly Adopts a Hoogsteen Conformation in the Absence of TnpA

We used NMR spectroscopy to characterize the solution conformation of the IS608 RE and LE hairpins in the absence of bound TnpA. For our studies, we used DNA constructs (Table S1 and Figure S1) containing the same sequences used to determine the crystal structures (PDB ID: 2VJV, 2A6O) of the IS608 TnpA-DNA complex^15^.

Remarkably, NMR analysis of the naked dinucleotide (TT) RE apical loop (Figure 2A), unambiguously showed that the A12-T9 bp forms a Hoogsteen conformation as the dominant state in solution even in the absence of TnpA protein. The chemical shift of the T9 imino proton at ∼13.25 ppm (Figure 2B) was upfield shifted relative to a Watson-Crick bp (∼13.65 ppm) and had a distance-based Nuclear Overhauser effect (NOE) contact with A12(H8) as expected for Hoogsteen but not Watson-Crick pairing (Figure 2C,D and Figure S3A-C)^20^. Several additional data establish that A12 adopts a *syn* conformation, including strong intra-nucleotide A12(H1ʹ)···A12(H8) and sequential A12(H8)···T11(H2ʹ/H2ʹʹ) NOEs (Figure 2D) as well as > 3 ppm downfield shifted A12(C8) and A12(C1ʹ) chemical shifts (Figure 2E, Figure S3D,E). The ∼2 ppm upfield shifted T9(N3) was also consistent with Hoogsteen pairing (Figure 2E). In addition to these Hoogsteen bp signatures, we also observed a long-range A12(H8)···T10(H2ʹ) NOE (Figure S4), which is to be expected based on the crystal structure of the IS608 TnpA-DNA complex and only for A12 in the syn but not the anti conformation. We did not observe the sequential A12(H2)···T11(H6) NOE, typically seen for Hoogsteen bps in duplex DNA^20^, but this is also to be expected given that T11 is part of the apical loop adopting an unusual conformation which lengthens the A12(H2)···T11(H6) distance to 6 Å instead of 4 Å in duplex DNA.

**Figure 2.**
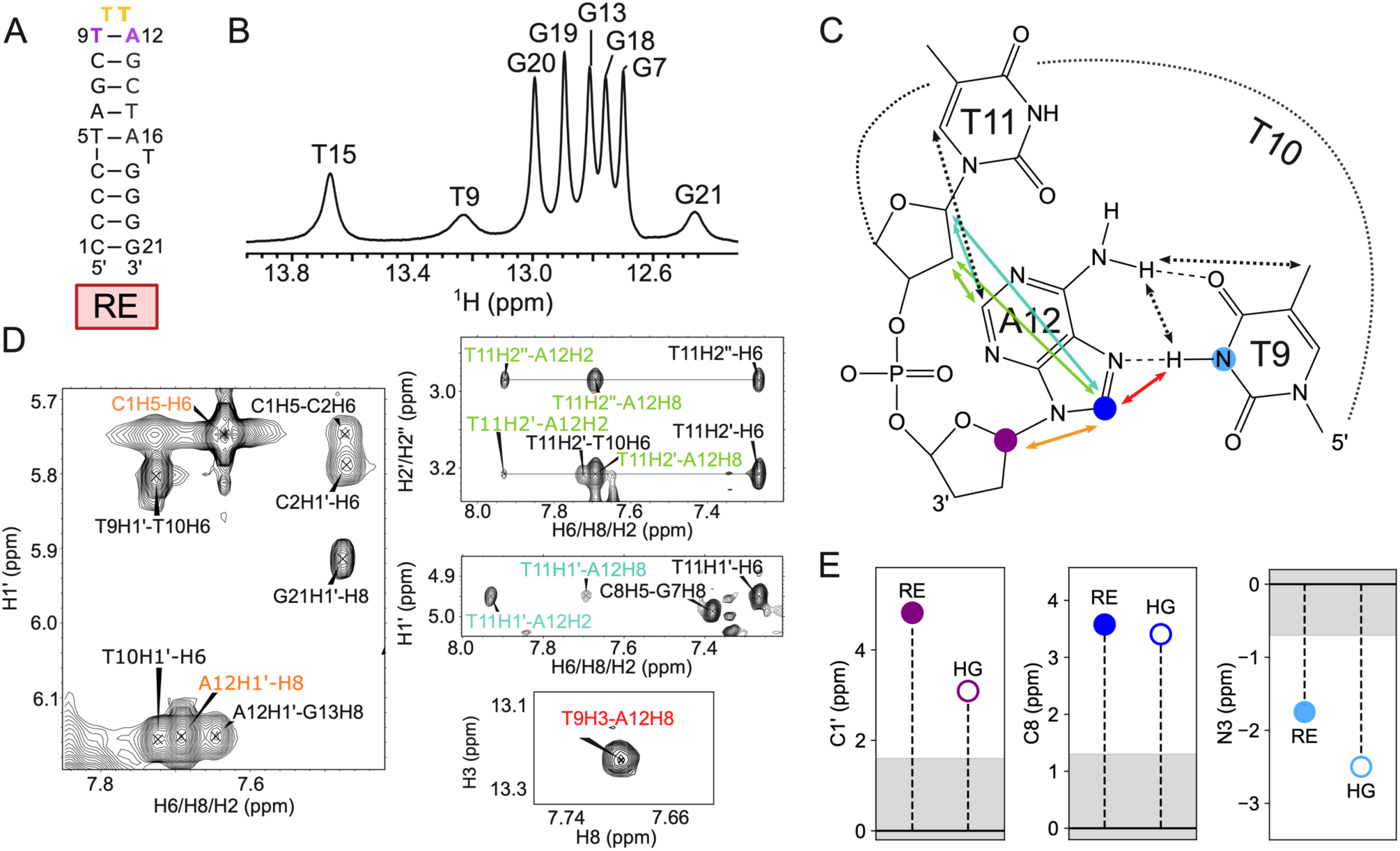
The A12-T9 base pair in RE forms a Hoogsteen conformation as the dominant state in the absence of TnpA. (A) The RE DNA construct used in this study was derived from the crystal structure of the TnpA-DNA complex (PDB ID: 2A6O). (B) 1D 1H NMR spectrum of RE showing imino resonances. The T9(H3) imino resonance belonging to the A12(T9) Hoogsteen base pair is upfield shifted as expected for Hoogsteen pairing^20^. (C) Chemical structure of the A12-T9 Hoogsteen base pair showing distance-based NOE contacts using color-coded arrows. The corresponding NOE cross-peaks are shown in D. The A(C1ʹ), A(C8), and T(N3) nuclei with NMR chemical shifts sensitive to the Hoogsteen versus Watson-Crick pairing are highlighted using colored circles and the chemical shift values are shown in E. (D) Regions of the 2D ^1^H-^1^H NOESY spectrum (mixing time 200 ms) showing the NOE cross-peaks and distant-based connectivity highlighted in C. The high intensity ratio of 0.95:1 between the NOE cross-peaks (orange) of A12(H1ʹ)···A12(H8) and the reference C1(H5)-C1(H6) indicates A12 is *syn*^9^. (E) Shown in filled circles are the differences between the chemical shifts measured for A(C1ʹ), A(C8), and T(N3) in the A12-T9 base pair of RE relative to the average chemical shifts for Watson-Crick base pairs derived from the BMRB^37^. For comparison, the difference between the chemical shifts typical of an A-T Hoogsteen base pair deduced using prior *R*_1ρ_ studies^38^ relative to the average chemical shifts for Watson-Crick base pairs derived from the BMRB^37^ are shown in open circles. The gray shaded area denotes the range of chemical shifts for the A-T Watson-Crick base pair derived from the BMRB^37^.

We also observed several additional NOEs (Table S2) including A12(H2)···T11(H2ʹ/H2ʹʹ), A12(H8)···T10(H1ʹ), A12(H8)···T11(H1ʹ), A12(H2)···T11(H1ʹ), T10(H6)···G13(H1ʹ), T10(H1ʹ)···G13(H1ʹ) and T10(H1ʹ)···C8(H1ʹ) involving apical loop residues which are consistent with the crystal structures^15, 17^. Indeed, all the NOEs predicted by the crystal structure were observed, with no extra NOEs detected (Figure S3A-C). Taken together, these results indicate that the unbound RE DNA hairpin is preorganized forming a conformation containing an A12-T9 Hoogsteen bp like that observed when bound to the TnpA protein.

### The A-T Base Pair in LE Predominantly Adopts a Watson-Crick Conformation in the Absence of TnpA

The LE construct (Figure 3A) differs from RE only by insertion of a single thymine residue (T*) in the apical loop, and crystal structures demonstrate that the A12-T9 bp also adopts a Hoogsteen conformation when the LE hairpin is bound to TnpA.

**Figure 3.**
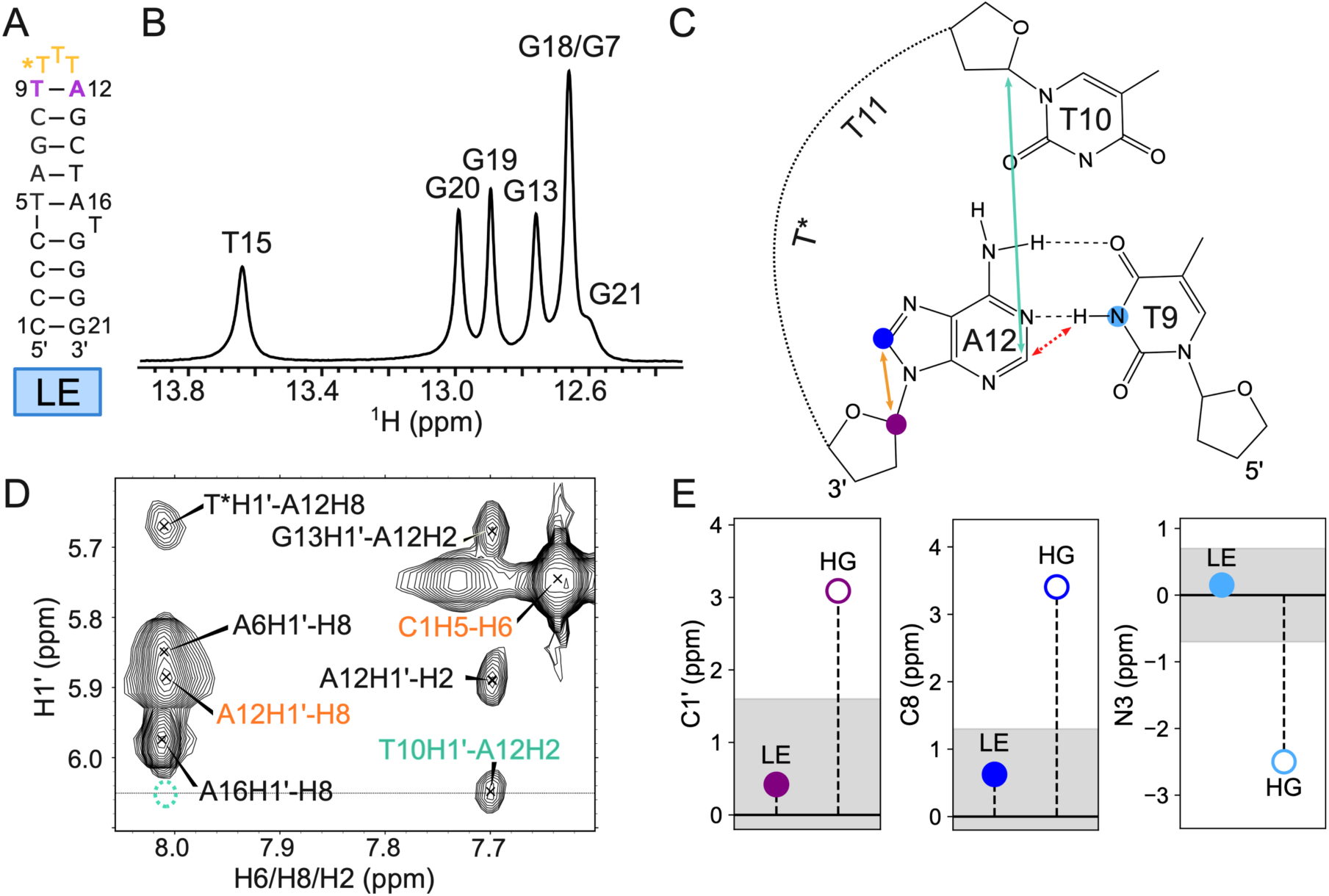
The A12-T9 base pair in LE forms a Watson-Crick conformation as the dominant state in the absence of TnpA. (A) The LE DNA construct used in this study was derived from the crystal structure of the TnpA-DNA complex (2VJV). (B) 1D ^1^H NMR spectrum of LE showing T(H3) imino resonances falling in the expected Watson-Crick base pair region. (C) Chemical structure of the A12-T9 Watson-Crick base pair showing distance-based NOE contacts using color-coded arrows. The corresponding NOE cross-peaks are shown in D. The A(C1ʹ), A(C8) and T(N3) nuclei with NMR chemical shifts sensitive to the Hoogsteen versus Watson-Crick conformation are highlighted in colored circles and the chemical shift values are shown in E. (D) Region of the 2D ^1^H-^1^H NOESY spectrum (mixing time 200 ms) showing the NOE cross-peaks and distant-based connectivity highlighted in C. The low-intensity ratio of 0.25:1 between the NOE cross-peaks (orange) of A12(H1ʹ)···A12(H8) and the reference C1(H5)-C(H6), together with the presence of the A12(H2)···T10(H1ʹ) cross-peak (cyan), indicates A12 is *anti*^9^. The dotted cyan circle denotes a lack of the A12(H8)···T10(H1ʹ) cross peak, also indicative of a Watson-Crick conformation. (E) Shown in filled circles are the differences between the chemical shifts measured for A(C1ʹ), A(C8), and T(N3) in the A12-T9 base pair of LE relative to the average chemical shifts for Watson-Crick base pairs derived from the BMRB^37^. For comparison, the difference between the chemical shifts typical of an A-T Hoogsteen base pair deduced using prior *R*_1ρ_ studies^38^ relative to the average chemical shifts for Watson-Crick base pairs derived from the BMRB^37^ are shown in open circles. The gray shaded area denotes the range of chemical shifts for the A-T Watson-Crick base pair derived from the BMRB^37^.

Strikingly, NMR analysis revealed that in stark contrast to RE, the A12-T9 bp in the trinucleotide LE construct does not adopt a Hoogsteen conformation in the absence of TnpA. Instead, the NMR data points to a Watson-Crick conformation in which A12 is *anti*. While the T9 imino proton was difficult to resolve in 1D ^1^H spectra due to resonance overlap (Figure 3B), it could be observed in a selectively ^13^C/^15^N labeled DNA sample (see below), and it had a chemical shift of 13.65 ppm consistent with an A-T Watson-Crick bp (Figure S5A). Several lines of evidence establish the A12 adopts the *anti* conformation, including a weak intra-nucleotide A12(H1ʹ)···A12(H8) NOE as well as A12(H2)···T10(H1ʹ) and A12(H2)···T10(H2ʹ/H2ʹʹ) NOEs (Figure 3C,D, Figure S5B,C, and Table S2). In addition, the A12(C8) and A12(C1ʹ) chemical shifts were no longer downfield shifted (Figure 3E and Figure S5D,E) consistent with A12 adopting the *anti* conformation. The sequential sugar(H1ʹ)-base(H6/8) NOEs were not interrupted at A12-T9, as expected for an intra-helical Watson-Crick conformation (Figure S5B,C and Table S2). Many of the NOEs observed in RE, including those characteristic of T11-T9 stacking and the T10-G13 contact, were notably absent in LE and there were new NOEs including A12(H2)···T10(H1ʹ/H2ʹ/H2ʹʹ) and A12(H2)···T*(H1ʹ/H2ʹ/H2ʹʹ), which were not observed in RE (Figure 3D and Table S2). Therefore, in contrast to RE, the LE apical loop is not preorganized for TnpA binding and the A12-T9 bp predominantly adopts a distinct Watson-Crick conformation.

### The A-T Base Pair Exists as a Dynamic Equilibrium Involving Hoogsteen and Watson-Crick Conformations

We hypothesized that in both LE and RE, the A12-T9 bp exists as a dynamic equilibrium between Hoogsteen and Watson-Crick conformations and that increasing the length of the apical loop preferentially stabilizes the Watson-Crick over the Hoogsteen conformation. To test this hypothesis, samples in which A12 and T9 residues were uniformly ^13^C/^15^N labeled. We then measured the micro-to-millisecond timescale dynamics of the A12-T9 bp using off-resonance ^13^C *R*_1ρ_ relaxation dispersion experiments^9, 21–22^ targeting A(C8), A(C2), A(C1ʹ), T(C1ʹ), and T(C6). We were unable to measure dispersion profiles for A12(C1ʹ) in LE due to spectral overlap.

Interestingly, both the A12(C1ʹ) and A12(C8) off-resonance *R*_1ρ_ profiles measured in RE (Figure 4A and Table S3) demonstrated that the A12-T9 Hoogsteen bp exists in dynamic equilibrium with a short-lived and lowly-populated conformational state. The *R*_1ρ_ profiles could be satisfactorily combined in a 2-state fit (Figure 4A, Figure S6A, and Table S4), revealing that the minor state has a population of ∼11 ± 1% and an exceptionally short lifetime of 55 ± 1 µs. The 2-state fit also revealed that the minor state has A12(C1ʹ) and A12(C8) chemical shifts (Table S4) that are substantially upfield shifted relative to the dominant ground-state (-4.6 ± 0.1 ppm and - 4.7 ± 0.1 ppm, respectively), falling in the region expected for a Watson-Crick conformation. Indeed, these chemical shifts were like those measured for the Watson-Crick A12-T9 bp in LE (Figure 3E and Figure 4B). Therefore, even though the dominant conformation of the A12-T9 bp in RE is Hoogsteen, the bp does form the Watson-Crick conformation as a minor state (Figure 4C). The minor Watson-Crick conformation also explains the absence of observable exchange for A12(C2), T9(C6) (Figure 4A), and T9(C1ʹ) (Figure S6B), as the chemical shifts of these nuclei have previously been shown to be insensitive to the Watson-Crick-Hoogsteen dynamics^9–10, 20, 23^.

**Figure 4.**
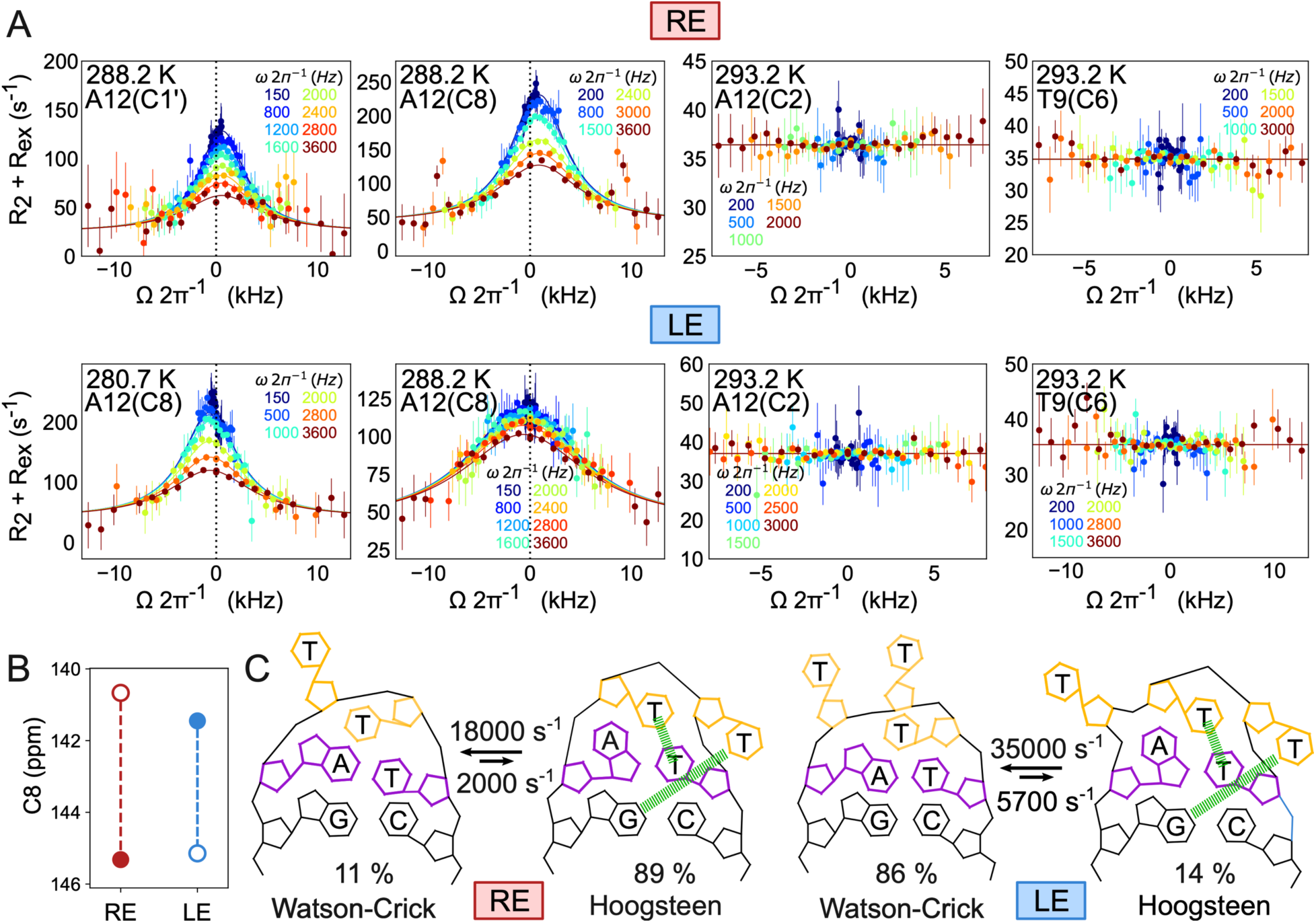
The A-T base pair exists as a dynamic equilibrium of Hoogsteen and Watson-Crick conformations. (A) Off-resonance ^13^C *R*_1ρ_ profiles measured for the A12-T9 base pair in RE (top) and on LE (bottom) at 900 MHz (600 MHz for A12(C1’)). The different spin-lock powers are color-coded. Shown is the fit (solid line) to a 2-state exchange model using the Bloch– McConnell equations, as described in Methods. Error bars represent the experimental uncertainty in the *R*_1ρ_ data and were obtained by propagating the error in *R*_1ρ_ using the Monte Carlo procedure. Dashed vertical line at 0 Hz offset shows how profiles are displaced along opposite directions for RE and LE, consistent with the inversion of the chemical shifts of the major and minor states. (B) Chemical shifts of A12(C8) in RE (red) and LE (blue) for the dominant (filled circles) and minor state (open circles). The chemical shift of the minor Watson-Crick state of RE is consistent with the chemical shift of the major Watson-Crick state of LE and *vice versa*. (C) NMR-derived conformational equilibria in RE and LE. The exchange parameters correspond to values measured at 15 °C.

The A12(C8) off-resonance *R*_1ρ_ profile measured in LE (Figure 4A and Table S5) also demonstrated that the A12-T9 Watson-Crick bp exists in dynamic equilibrium with a short-lived and lowly-populated conformational state (Figure 4A and Figure S6C). Because the exchange rate was slightly faster in the trinucleotide LE relative to the dinucleotide RE, we performed additional *R*_1ρ_ measurements at a lower temperature of 7.5 °C to better define the exchange parameters (Figure 4A). A 2-state fit of these data revealed a minor state population of ∼14 ± 2% and an exceptionally short lifetime of 28 ± 1 µs. Interestingly, the A12(C8) chemical shift of the minor state was downfield shifted by 3.6 ± 0.2 ppm relative to the dominant state, falling in the region expected for a Hoogsteen conformation (Figure 4B, Figure S6C, and Table S6) and like the chemical shift measured for the Hoogsteen A12-T9 bp in RE (Figure 2E). Therefore, even though the dominant conformation of the A12-T9 bp in LE is Watson-Crick, the Hoogsteen conformation forms as a minor state (Figure 4C and Table S6) with a population of 14%, which is an order of magnitude greater than the ∼1% population typical of canonical duplex DNA^9–10^. Once again, Hoogsteen as the minor conformation explains the lack of chemical exchange at A12(C2), T9(C6) (Figure 4B), and T9(C1’) (Figure S6B).

Taken together, these results show that the A12-T9 bp exists in a dynamic equilibrium between the Hoogsteen and Watson-Crick conformations and that increasing the apical loop length from two to three nucleotides inverts the equilibrium rendering Watson-Crick the dominant state and Hoogsteen the minor state (Figure 4B,C). In both RE and LE, the fast exchange kinetics explains the appearance of a single set of resonances despite the relatively large population of the minor state. Below, we leverage these motionally averaged chemical shifts to assess the interactions responsible for stabilizing the Hoogsteen conformation.

### The Relative Stability of the Hoogsteen Versus Watson-Crick Conformation Depends on the Identity of Apical Loop Residues

Crystal structures of RE and LE hairpins in complex with TnpA reveal several intra-molecular DNA contacts involving the thymine apical loop residues (Figure 1D). These contacts may stabilize the Hoogsteen conformation directly through stacking with T11 or indirectly via hydrogen bonding between T10 and G13 (Figure 1D). To test the importance of these interactions, we prepared RE samples (Ab10 and Ab11, Figure 5A) in which either T10 or T11 was substituted with an abasic site lacking the base all together. NMR analysis (Figure S7A,B) revealed that in both cases, the A12-T9 bp conformation shifted toward Watson-Crick, with a more pronounced shift observed for Ab10 (Figure 5B,C), highlighting the critical role of the G13(N3)-T10(H3) hydrogen bond in stabilizing the A12-T9 Hoogsteen conformation.

**Figure 5.**
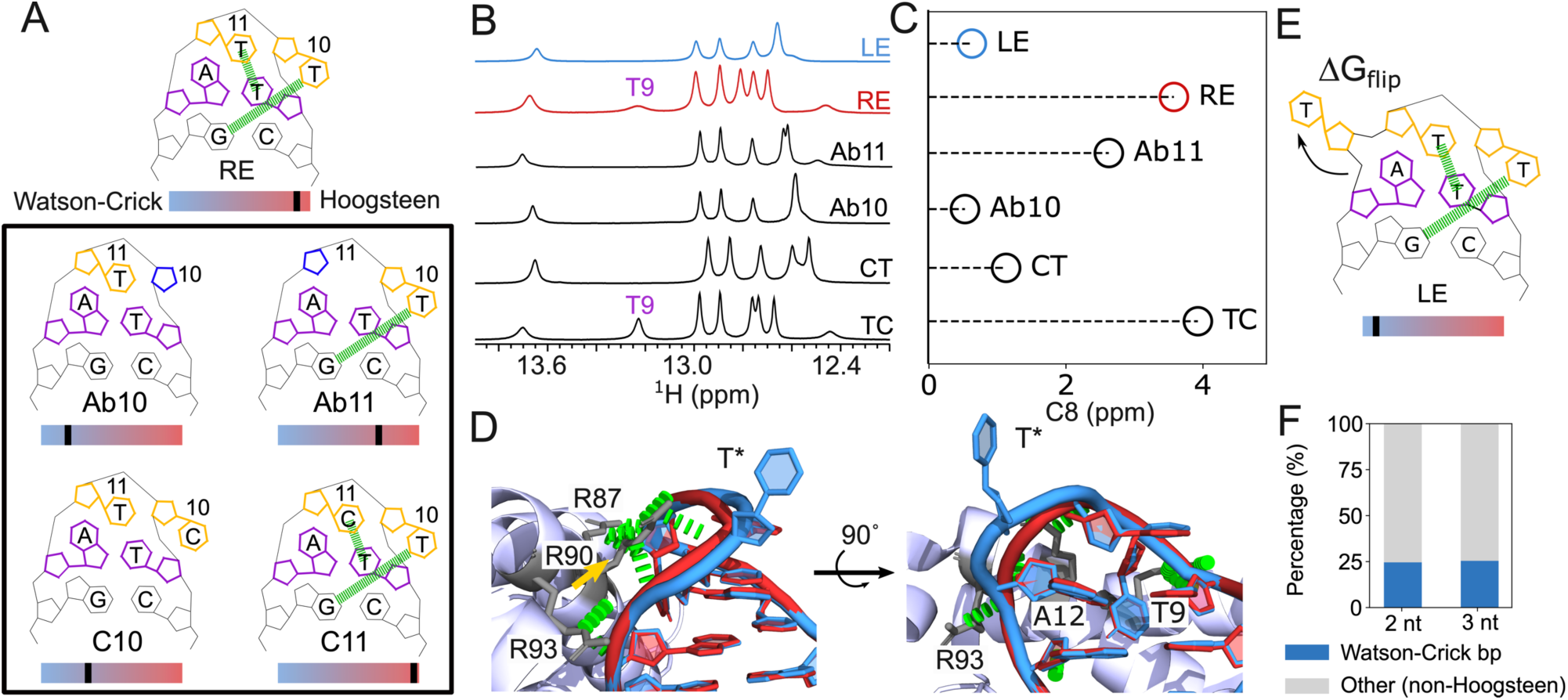
Hoogsteen propensity depends on the identity of apical loop residues. (A) Schematic showing abasic (Ab) substitutions (Ab10 and A11) and mutations (C10 and C11) designed to impact interaction shown in green on wt RE. Comparison of (B) 1D ^1^H NMR spectra and (C) A(C8) chemical shifts across the different variants shown in (A) to assess Hoogsteen versus Watson-Crick pairing. Shown are the differences between the observed A12(C8) chemical shifts and the average chemical shift for an A-T Watson-Crick base pair obtained from the BMRB^37^. (D) Comparison of the RE (red) and LE (blue) apical loops and their adjacent base pairs within the binding pocket of the TnpA transposase. Contacts between TnpA arginine and the DNA phosphate groups are shown in green. (E) The A12(*syn*)-T9 conformation in LE bound to TnpA is achieved by extra-helical flipping of the inserted T* residue while the remaining residues adopt a conformation like that of the dinucleotide apical loop. (F) Survey of apical loop structures in RNA (see methods). Shown is the percentage of A-U base pairs which form Watson-Crick or other conformations in dinucleotide (2 nt) and trinucleotide (3 nt) apical loops.

For the ISDra2 sequence family, both RE and LE tend to have dinucleotide apical loops^14^ and there is a naturally occurring LE in which the TT apical loop motif is replaced with TC. A crystal structure of this hairpin bound to ISDra2 TnpA (PDB ID: 2XQC) shows a conformation nearly identical to that of the TT apical loop with A12-T9 adopting a Hoogsteen conformation and in which C11 instead of T11 stacks on T9. Indeed, NMR analysis (Figure S7C) revealed that substituting T11 with C11 resulted in the A12-T9 bp maintaining a Hoogsteen conformation (Figure 5B,C).

Interestingly, in contrast to TC, there are no naturally occurring CT dinucleotide sequences (Figure S8). Moreover, such a T10 to C10 substitution is predicted to disrupt the G13(N3)-T10(H3) hydrogen bond, which we have shown is crucial for maintaining the A12-T9 bp in a Hoogsteen conformation (Figure 5A). Indeed, there is a crystal structure (PDB ID: 4R8U)^24^ for an A-T bp adjacent to a CT dinucleotide apical loop derived from a different system and in this case it adopts a Watson-Crick conformation. As expected, our NMR analysis (Figure S7D) also showed that substituting T10 with C10 resulted in the A12-T9 bp predominantly adopting a Watson-Crick conformation (Figure 5B,C). The importance of the G13(N3)-T10(H3) hydrogen bond in stabilizing the A12-T9 Hoogsteen conformation is also highlighted by another crystal structure (PDB ID: 4ER8) in which the G13-C8 was flipped to C13-G8 thus disrupting the hydrogen bond. As expected, the A12-T9 bp in this structure adopted a Watson-Crick conformation.

Taken together, these results show that the Hoogsteen bp is stabilized by specific interactions between apical loop residues and adjacent bps as well as define 5’-CTT(T/C)AG-3’ as a Hoogsteen-specific apical loop motif.

### The Interplay Between Shape-Recognition and Conformational Penalty Dictates Hoogsteen Propensity in the Trinucleotide Apical Loop

Why does the A12-T9 bp in the naked trinucleotide LE apical loop predominantly adopt a Watson-Crick conformation and what drives it to adopt a Hoogsteen conformation when bound to TnpA? TnpA recognizes the DNA hairpins through several shape-specific contacts with the DNA backbone, including several salt bridges involving arginine residues and DNA backbone phosphate groups (Figure 5D). Many of these contacts depend on the A12-T9 bp adopting a Hoogsteen conformation. Attempts to model an A12-T9 Watson-Crick bp resulted in the loss of some of these contacts and steric clashes (Figure S9A). Furthermore, when predicting 3D structures for TnpA-LE complexes using AlphaFold3 (AF3)^25^, half of the predicted structures had the A12-T9 bp in a Hoogsteen conformation and none were canonical Watson-Crick bps (Figure S9B). Interestingly, in many of the AF3 predicted structures, the A12(*anti*)-T9 bp was constricted with C1ʹ-C1ʹ distance < 10 Å (Figure S9B). This result indicated that constriction at the A12-T9 bp, facilitated by the Hoogsteen conformation, is likely critical for forming the specific configuration recognized by TnpA (Figure S9B).

To bind TnpA and form these Hoogsteen-dependent contacts, the LE hairpin robustly adopts a conformation containing an A12-T9 Hoogsteen bp nearly identical to that of the dinucleotide RE hairpin as revealed in several crystal structures (Figure 5D and Figure S1). The heavy atom RMSD over apical loop residues is ∼0.6 Å. This mimicry is achieved through extra-helical flipping of the additional T* residue, which enables the remaining residues to adopt a conformation like that of RE (Figure 5E). Extra-helical flipping of the T* residue also avoids steric clashes with R90 in TnpA (Figure 5D). Thus, through shape-specific contacts with a Hoogsteen-dependent shape, TnpA renders Hoogsteen the dominant conformation of the A-T bp in the trinucleotide LE hairpin without directly interacting with the Hoogsteen bp itself (Figure 5D). Because this Hoogsteen conformation already has a sizeable population of ∼14% in naked LE (Figure 4C), a small energetic bias of ∼ 2 kcal/mol would be sufficient to invert the Watson-Crick to Hoogsteen equilibrium.

The requirement to flip out the T* apical loop residue to accommodate the A12-T9 Hoogsteen bp may also explain why, in the absence of TnpA, the dominant conformation in LE is Watson-Crick not Hoogsteen (Figure 5E). In single-stranded RNA, the extra-helical flipping of nucleotides incurs an energetic cost of ∼1.5–2.0 kcal/mol^26^ and a similar penalty is likely applicable to single-stranded DNA. This additional energetic cost incurred by LE relative to RE is comparable to the ∼2 kcal/mol energetic bias required to invert the Watson-Crick-to-Hoogsteen equilibrium, and it might explain the reduced propensity of the A12-T9 bp to adopt the Hoogsteen conformation in LE compared to RE.

### No A-U Hoogsteen bps Observed in Corresponding RNA Apical Loops

Given the more extensive database of RNA structures in the PDB relative to DNA as well as RNA’s tendency to adopt a wide variety of non-canonical conformations, we wondered whether the Hoogsteen conformation observed near the apical loop is unique to DNA or if similar conformations could also occur in RNA. This question is particularly interesting given prior studies showing that Hoogsteen bps are strongly disfavored in A-form RNA compared to B-form DNA^27–28^. This energetic instability has been attributed to steric clashes between the *syn* purine base and the RNA backbone, as well as the higher energetic cost of bringing bases into closer proximity to form hydrogen bonds^27–28^.

To examine this question, we surveyed existing crystal structures of RNA for A-U bps forming adjacent to di- and trinucleotide apical loops (Methods). Strikingly, among 63 dinucleotide RNA apical loops from 49 different crystal structures (Table S7), none of the A-U bps were Hoogsteen (Figure 5F). Moreover, in all cases, the adenine base was *anti* rather than *syn*. Approximately 25% of the bps were Watson-Crick, while the remaining 75% exhibited non-canonical conformations (Figure 5F and Table S7). Similarly, among 228 trinucleotide RNA apical loops with an adjacent A-U bp from a total of 144 crystal structures (Table S8), none were Hoogsteen (Figure 5F). Only one bp had adenine in a *syn* conformation, but the C1ʹ-C1ʹ distance was not constricted and the bp did not form Hoogsteen-type hydrogen bonds (PDB ID: 6N5N). Like RNA hairpins with dinucleotide apical loops, 25% of the bps were Watson-Crick while 75% adopted non-canonical conformations (Figure 5F and Table S8).

These results indicate that the A-T Hoogsteen bp forming adjacent to the dinucleotide apical loop is unique to DNA and does not form with comparable propensity in RNA apical loops.

## Conclusion

Hairpins capped by apical loops forming in single-stranded DNA regions during replication and transcription are implicated in a variety of processes including horizontal gene transfer^13, 29^, recombination^11, 18^, regulation of gene expression^30^, replication^31–32^, DNA damage repair^33^, and genome stability maintenance^34–36^. Despite their functional significance, structural information on DNA apical loops remains surprisingly limited. Here, we demonstrated that even in the absence of TnpA, the 5’-CTT(T/C)AG-3’ apical loop motif adopts a unique conformation stabilized by an A-T Hoogsteen closing bp. This finding shows that functionally relevant Hoogsteen bps, which are specifically recognized by proteins, can exist as the dominant state in naked DNA. Moreover, the propensity to adopt the A-T Hoogsteen conformation was tunable, depending on the length and composition of the apical loop. Further studies are required to determine whether the different propensities to form Hoogsteen bps in LE versus RE shape TnpA binding at these sites and impact transposition efficiency. Notably, Hoogsteen bps were not observed in RNA apical loops, potentially offering a new basis for distinguishing DNA hairpins from their RNA counterparts. We anticipate that other families of DNA apical loop motifs featuring A(*syn*)-T and potentially also G(*syn*)-C^+^ Hoogsteen bps remain to be identified that may play broad and yet to be characterized roles in DNA structure and function.

## Acknowledgments

We would like to thank members of the Al-Hashimi lab for their input. HMA is a member of the New York Structure Biology Center (NYSBC). This work was supported by the NIH National Institute of General Medical Sciences (R01GM089846). The data collected at NYSBC using the 900 MHz spectrometers were purchased with funds from NIH grant P41GM066354 and the New York State Assembly. Data collected using the 900MHz NEO spectrometer is supported by NIH grant S10OD030373. Some of the work presented here was conducted at the Center on Macromolecular Dynamics by NMR Spectroscopy located at NYSBC, supported by a grant from the NIH National Institute of General Medical Sciences RM1GM145397. OS is an Awardee of the Women’s Postdoctoral Career Development Award from the Weizmann Institute of Science, and the Zuckerman-CHE Israeli Women Postdoctoral Program Fellowship.

## Author contributions

Conceptualization: SG1, HMA, and OS. PDB structure analysis: SG1 and KS. NMR measurements and analysis: SG1, OS, SG2 and SP. RNA structural survey: AG. Funding acquisition: HMA. Supervision: HMA. Writing original draft: HMA, SG1, OS.

## Supplementary Materials

Materials and Methods

Figures S1 to S9

Tables S1 to S8

## References

1. Watson, J. D.; Crick, F. H., Molecular structure of nucleic acids; a structure for deoxyribose nucleic acid. Nature 1953, 171 (4356), 737–8.

2. Hoogsteen, K., Unit-cell dimensions and space group of 6-mercaptopurine monohydrate. Nature 1956, 178 (4529), 379.

3. Abrescia, N. G., et al., Crystal structure of an antiparallel DNA fragment with Hoogsteen base pairing. Proceedings of the National Academy of Sciences of the United States of America 2002, 99 (5), 2806–11.

4. Kitayner, M., et al., Diversity in DNA recognition by p53 revealed by crystal structures with Hoogsteen base pairs. Nature structural & molecular biology 2010, 17 (4), 423–9.

5. Nair, D. T., et al., Replication by human DNA polymerase-iota occurs by Hoogsteen base-pairing. Nature 2004, 430 (6997), 377–80.

6. Shi, H., et al., Revealing A-T and G-C Hoogsteen base pairs in stressed protein-bound duplex DNA. Nucleic Acids Res 2021, 49 (21), 12540–12555.

7. Park, J. Y.; Choi, B. S., NMR investigation of echinomycin binding to d(ACGTTAACGT)2: Hoogsteen versus Watson-Crick A.T base pairing between echinomycin binding sites. J Biochem 1995, 118 (5), 989–95.

8. Nikolova, E. N., et al., Probing transient Hoogsteen hydrogen bonds in canonical duplex DNA using NMR relaxation dispersion and single-atom substitution. J Am Chem Soc 2012, 134 (8), 3667–70.

9. Nikolova, E. N., et al., Transient Hoogsteen base pairs in canonical duplex DNA. Nature 2011, 470 (7335), 498–502.

10. Nikolova, E. N., et al., Guanine to inosine substitution leads to large increases in the population of a transient G.C Hoogsteen base pair. Biochemistry 2014, 53 (46), 7145–7.

11. Bikard, D., et al., Folded DNA in action: hairpin formation and biological functions in prokaryotes. Microbiol Mol Biol Rev 2010, 74 (4), 570–88.

12. Guynet, C., et al., In vitro reconstitution of a single-stranded transposition mechanism of IS608. Mol Cell 2008, 29 (3), 302–12.

13. Ton-Hoang, B., et al., Transposition of ISHp608, member of an unusual family of bacterial insertion sequences. EMBO J 2005, 24 (18), 3325–38.

14. Pasternak, C., et al., Irradiation-induced Deinococcus radiodurans genome fragmentation triggers transposition of a single resident insertion sequence. PLoS Genet 2010, 6 (1), e1000799.

15. Barabas, O., et al., Mechanism of IS200/IS605 family DNA transposases: activation and transposon-directed target site selection. Cell 2008, 132 (2), 208–20.

16. Hickman, A. B., et al., DNA recognition and the precleavage state during single-stranded DNA transposition in D. radiodurans. EMBO J 2010, 29 (22), 3840–52.

17. Morero, N. R., et al., Targeting IS608 transposon integration to highly specific sequences by structure-based transposon engineering. Nucleic Acids Res 2018, 46 (8), 4152–4163.

18. Kersulyte, D., et al., Transposable element ISHp608 of Helicobacter pylori: nonrandom geographic distribution, functional organization, and insertion specificity. J Bacteriol 2002, 184 (4), 992–1002.

19. Meyder, A., et al., Estimating Electron Density Support for Individual Atoms and Molecular Fragments in X-ray Structures. J Chem Inf Model 2017, 57 (10), 2437–2447.

20. Sathyamoorthy, B., et al., Insights into Watson-Crick/Hoogsteen breathing dynamics and damage repair from the solution structure and dynamic ensemble of DNA duplexes containing m1A. Nucleic acids research 2017, 45 (9), 5586–5601.

21. Koss, H., et al., Site-based description of R(1)(rho) relaxation in local reference frames. J Magn Reson 2023, 347, 107366.

22. Rangadurai, A., et al., Characterizing micro-to-millisecond chemical exchange in nucleic acids using off-resonance R(1rho) relaxation dispersion. Prog Nucl Magn Reson Spectrosc 2019, 112-113, 55–102.

23. Shi, H., et al., Atomic structures of excited state A-T Hoogsteen base pairs in duplex DNA by combining NMR relaxation dispersion, mutagenesis, and chemical shift calculations. J Biomol NMR 2018, 70 (4), 229–244.

24. Zhou, H., et al., Characterizing Watson-Crick versus Hoogsteen Base Pairing in a DNA-Protein Complex Using Nuclear Magnetic Resonance and Site-Specifically (13)C- and (15)N-Labeled DNA. Biochemistry 2019, 58 (15), 1963–1974.

25. Abramson, J., et al., Accurate structure prediction of biomolecular interactions with AlphaFold 3. Nature 2024, 630 (8016), 493–500.

26. Sadee, C., et al., A comprehensive thermodynamic model for RNA binding by the Saccharomyces cerevisiae Pumilio protein PUF4. Nat Commun 2022, 13 (1), 4522.

27. Rangadurai, A., et al., Why are Hoogsteen base pairs energetically disfavored in A-RNA compared to B-DNA? Nucleic Acids Res 2018, 46 (20), 11099–11114.

28. Zhou, H., et al., m(1)A and m(1)G disrupt A-RNA structure through the intrinsic instability of Hoogsteen base pairs. Nat Struct Mol Biol 2016, 23 (9), 803–10.

29. Gonzalez-Perez, B., et al., Analysis of DNA processing reactions in bacterial conjugation by using suicide oligonucleotides. EMBO J 2007, 26 (16), 3847–57.

30. Hatfield, G. W.; Benham, C. J., DNA topology-mediated control of global gene expression in Escherichia coli. Annu Rev Genet 2002, 36, 175–203.

31. Heller, R. C.; Marians, K. J., Replication fork reactivation downstream of a blocked nascent leading strand. Nature 2006, 439 (7076), 557–62.

32. Hacker, K. J.; Alberts, B. M., The rapid dissociation of the T4 DNA polymerase holoenzyme when stopped by a DNA hairpin helix. A model for polymerase release following the termination of each Okazaki fragment. J Biol Chem 1994, 269 (39), 24221–8.

33. Trinh, T. Q.; Sinden, R. R., Preferential DNA secondary structure mutagenesis in the lagging strand of replication in E. coli. Nature 1991, 352 (6335), 544–7.

34. Gordenin, D. A., et al., Inverted DNA repeats: a source of eukaryotic genomic instability. Mol Cell Biol 1993, 13 (9), 5315–22.

35. Lewis, S. M., P nucleotide insertions and the resolution of hairpin DNA structures in mammalian cells. Proc Natl Acad Sci U S A 1994, 91 (4), 1332–6.

36. Sinden, R. R., et al., On the deletion of inverted repeated DNA in Escherichia coli: effects of length, thermal stability, and cruciform formation in vivo. Genetics 1991, 129 (4), 991–1005.

37. Hoch, J. C., et al., Biological Magnetic Resonance Data Bank. Nucleic Acids Res 2023, 51 (D1), D368–D376.

38. Alvey, H. S., et al., Widespread transient Hoogsteen base pairs in canonical duplex DNA with variable energetics. Nat Commun 2014, 5, 4786.

