## Supporting Methods, Figures and Tables for "An A-T Hoogsteen base pair in a naked DNA hairpin motif: A Protein-Recognized Conformation"

**This file includes:**

Materials and Methods

Supplementary Figures S1 to S9

Supplementary Tables S1 to S8

Supplementary References

### Materials and Methods

#### Inspection of crystal structures

To assess potential modeling errors in the A12-T9 base pairs (bps), we manually examined the electron density maps and structural models for all IS608 (PDB IDs: 6FI8, 2VJV, 2VIC, 2VHG, 2VIH, 2VJU, 2A6O) and ISDra2 (PDB IDs: 2XMA, 2XM3, 2XO6, 2XQC) structures deposited in the PDB. Two structural models were generated for each bp using the procedure developed by Shi et al[1] using the *PHENIX* software[2]: one in a Watson-Crick conformation and the other in Hoogsteen. The models were then compared by analyzing the correlation with the $F_c$ and $2F_o$–$F_c$ electron density maps. Steric clashes were assessed using MolProbity[3].

#### Sample preparation

<u>Synthesis and purification of DNA oligonucleotides.</u> Unlabeled single-stranded DNA oligonucleotides (~ 2 mg) corresponding to RE, LE, Ab10, Ab11, TC and CT (Table S1) were purchased from Integrated DNA Technologies with standard desalting purification. Selectively $^{13}C/^{15}N$-site-labeled single-stranded DNA oligonucleotides for RE and LE (Table S1) were purchased from Yale Keck Oligonucleotide Synthesis Facility with cartridge purification and using 2′-deoxyadenosine ($^{13}C$, 98%; $^{15}N$, 98%) and 2′-deoxythymidine ($^{13}C$, 98%; $^{15}N$, 98%) phosphoramidites obtained from Cambridge Isotope Laboratories.

<u>NMR Buffer.</u> The buffer used in all NMR experiments consisted of 15 mM sodium phosphate, 25 mM sodium chloride and 0.1 mM ethylenediaminetetraacetic acid (EDTA) in 90% $H_2O$:10%$D_2O$ at pH 6.8

<u>Preparation of NMR samples.</u> DNA hairpin samples were prepared by heating the single-stranded DNA oligonucleotides (~ 50 μM) to 95 °C for 5 min in water followed by fast cooling on ice for ~

30 mins. The sample was then buffer exchanged into NMR buffer at 4 °C using Amicon Ultra-15 centrifugal concentrators (3-kDa cutoff, Millipore Sigma) to a final nucleic acid concentration of ~ 1 mM. Extinction coefficients used to measure the oligonucleotide sample concentrations were estimated using the ADT Bio Oligo Calculator (https://www.atdbio.com/tools/oligo-calculator). Oligonucleotide sample concentrations ranged between 0.75 and 1.50 mM.

**NMR experiments**

Resonance assignment. 2D $^{1}H$-$^{13}C$ HSQC , $^{1}H$-$^{15}N$ SOFAST-HMQC[4] and $^{1}H$-$^{1}H$ NOESY were collected on a 600 MHz Bruker Avance-III HD spectrometer equipped with a 5 mm cryoprobe (TCI 600 H&F-C/N-D-5-Z) or on 800 MHz Bruker Avance spectrometer equipped with a 5 mm cryoprobe (TXO 800 H-C/N-D-05-Z) using Topspin 3.2 at T=10 °C. Data were processed and analyzed with NMRpipe[5] and SPARKY[6].

$^{13}C$ $R_{1\rho}$ relaxation dispersion (RD). Off-resonance $^{13}C$ $R_{1\rho}$ RD experiments[7-9] were performed on a Bruker Avance Neo 900 MHz spectrometer equipped with a 5-mm triple-resonance cryogenic probe (TCI 900 H&F-C/N-D-5-Z), or a 600 MHz Bruker Avance-III HD spectrometer equipped with a 5 mm cryoprobe (TCI 600 H&F-C/N-D-5-Z), and implemented using a 1D selective excitation scheme as described in prior studies[5, 7]. The spin-lock powers ($\omega_1/2\pi$) ranged from 150 to 5000 Hz, while the offsets ranged from ±3.5 times the spin-lock power (Table S2 and Table S4). For each resonance, 4-5 delay times were used during the relaxation period with a maximum duration between 8 and 32 ms.

Analysis of $R_{1\rho}$ data. The $R_{1\rho}$ data was analyzed as described previously[10]. Peak intensities were extracted using NMRPipe[5], and fitted to a mono-exponential decay as a function of the delay time to obtain the $R_{1\rho}$ value for the different spin-lock power and offset combinations. The error in $R_{1\rho}$ was estimated using a Monte Carlo procedure as described previously[11]. $R_{1\rho}$ values, measured as a function of different spin-lock power and offset combinations, were fit to a two-state exchange model using the Bloch-McConnell equations[12] to extract exchange parameters including the population of the minor state ($p_{minor}$), the exchange rate between the minor and dominant state ($k_{ex} = k_{forward} + k_{reverse}$), the difference between the chemical shifts of the minor and dominant states ($\Delta\omega = \omega_{minor} - \omega_{dominant}$), as well as transverse ($R_2$) and longitudinal ($R_1$) relaxation rates. In all cases, it was assumed that $R_{2,dominant}=R_{2,minor}=R_2$ and $R_{1,dominant}=R_{1,minor}=R_1$. The initial alignment of magnetization during the Bloch-McConnell simulations was determined based on the $k_{ex}/\Delta\omega$ ratio, as described previously[11]. The uncertainty in the exchange parameters was obtained using a Monte-Carlo scheme as described previously[13]. The fitted exchange parameters are summarized in Table S4 and Table S6. Global fitting of the $R_{1\rho}$ data was performed by sharing $k_{ex}$ and $p_{minor}$ in RE for A12(C1') and A12(C8). Since the LE A12(C8) resonance did not show a detectable chemical shift difference with a decrease of the temperature down to 7.5 °C, a global fit of the A12(C8) $R_{1\rho}$ data was performed across four temperatures sharing Δω, yielding exchange parameters within error of the those measured at 15 °C. Off-resonance $R_{1\rho}$ profiles (Figure 4A, Figure S6) were generated by plotting $(R_2 + R_{ex}) = (R_{1\rho} - R_1\cos^2\theta)/\sin^2\theta$, in which θ is the angle between the effective field of the observed resonance and the z-axis, as a function of $\Omega_{eff} = \omega_{obs} - \omega_{RF}$, in which $\omega_{obs}$ is the Larmor frequency of the observed resonance and $\omega_{RF}$ is the angular frequency of the applied spin-lock. Errors in $(R_2 + R_{ex})$ were determined by propagating the error in $R_{1\rho}$ obtained as described above.

**Modeling Watson-Crick the conformation into structures of the DNA-TnpA complex**

We modeled an A12-T9 Watson-Crick bp into the structure of the DNA-TnpA complex using the structure of LE bound to TnpA (PDB ID: 2VIC) as a template. To model the Watson-Crick bp, we utilized a DNA hairpin structure (PDB ID: 1UUT) containing a trinucleotide apical loop and an adjacent Watson-Crick A-T bp. This structure comprised the three-nucleotide apical loop, an adjacent Watson-Crick A-T bp, and the three Watson-Crick bps in the stem. PyMOL (The PyMOL Molecular Graphics System, Version 3.0, Schrödinger, LLC) was used to superimpose the three stem bps onto corresponding bps in the LE hairpin from the TnpA-DNA complex (PDB ID: 2VIC). The resulting structural model was analyzed to assess the preservation of TnpA-DNA interactions and identify any potential steric clashes between TnpA and the DNA.

Additionally, a series of TnpA-LE complex structures were predicted using AlphaFold3[14], utilizing the same protein sequences as in PDB ID: 2VIC and the LE sequence used in the NMR experiments (Table S1). A total of 20 structures were generated. In models containing A12(*anti*)-T bps, the C1'–C1' distance was reduced to <10 Å, classifying these bps as non-canonical Watson-Crick-like conformations.

**RNA hairpin PDB survey**

We conducted a survey of existing RNA crystal structures, focusing on A-U bps adjacent to dinucleotide and trinucleotide apical loops. X-ray structures with resolutions of ≤3.0 Å containing nucleic acids were downloaded from the RCSB Protein Data Bank (PDB)[15] on January 11, 2024. We processed the retrieved 11,133 PDB structures using X3DNA-DSSR[16] to create a searchable structural database. Structures identified by X3DNA as RNA hairpins were selected for further analysis. To identify 5′A-U3′ and 5′U-A3′ RNA bps adjacent to di- and trinucleotide apical loops, we defined them as hydrogen-bonded bases on the same RNA strand, separated by 2 or 3 nucleotides. The A-U bps were classified based on three criteria: (1) the conformation of the adenine (*syn*, *anti*, or non-canonical, denoted as "---" in X3DNA); (2) the C1'–C1' distance; and (3) the hydrogen-bonding pattern. The Watson-Crick conformation was defined by having adenine in the *anti* conformation, a C1'–C1' distance between 10.0 Å and 10.8 Å, and the bases stabilized by two hydrogen bonds: N3(imino)-N1 and O4(carbonyl)-N6(amino). The Hoogsteen conformation was defined by having an adenine in the *syn* conformation, the C1'–C1' distance

≤10 Å, and the bases stabilized by at least two hydrogen bonds: N3(imino)-N7 and O4(carbonyl)-N6(amino). Other conformations were classified as "other", including: Hoogsteen-like conformations with C1'–C1' distance >10 Å, *trans*-Hoogsteen bps with adenine in the *anti* conformation and only one hydrogen bond, and bps identified by X3DNA as *trans* Hoogsteen/Watson-Crick (tHW), *trans* Sugar/Hoogsteen (tSH), *cis* Sugar/Hoogsteen (cSH), or *cis* Sugar/Watson-Crick (cSW)[17].

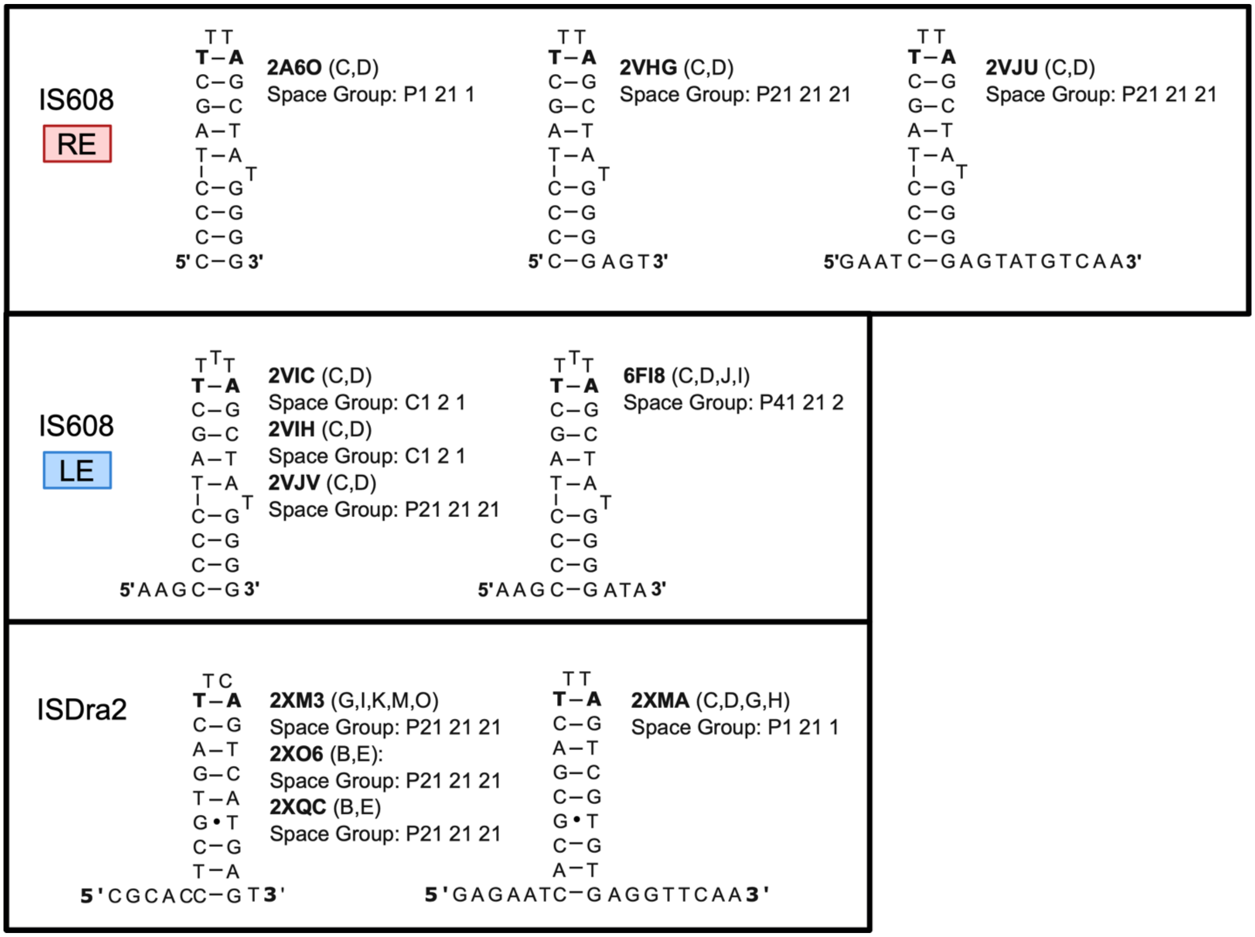


**Figure S1.** DNA constructs used in crystal structures of the TnpA-DNA complex for RE (top) and LE (middle) of IS608 and ISDra2 (bottom, left is LE and right is RE) from the IS200/IS605 family. In all the structures, the A-T base pair adjacent to the apical loop (indicated using bold letters) adopted a Hoogsteen conformation.

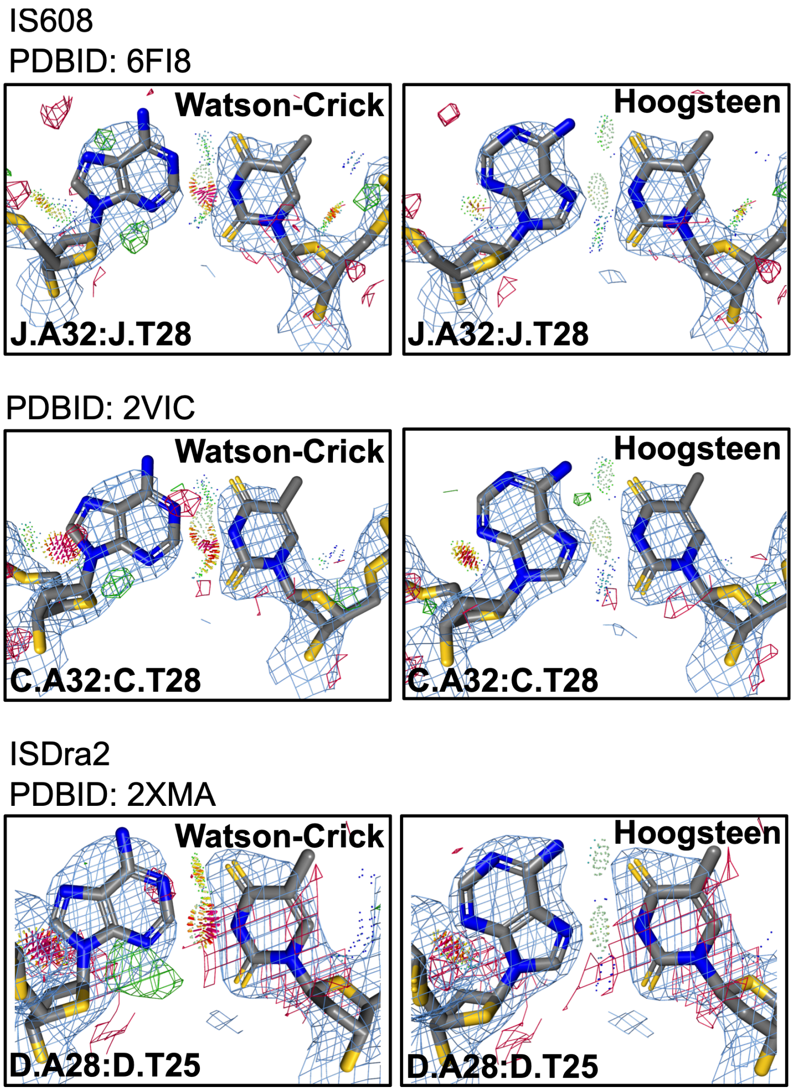


**Figure S2. Hoogsteen and Watson-Crick models for IS608 and ISDra2 DNA hairpins in complex with TnpA.** Comparison of the original Hoogsteen (right) and corresponding mismodeled Watson-Crick (left) models for A-T base pairs adjacent to the apical loop in IS608 and ISDra2 hairpins. Gray and purple meshed regions represent $2mF_o$-$DF_c$ densities at 1.0σ and 3.0σ, respectively, while blue and red meshed regions are $mF_o$-$DF_c$ difference densities contoured at 3.0σ and -3.0σ, respectively. Also shown is the stereochemistry assessed by MolProbity (see Methods). In all cases the agreement between the chemical model and experimental data, together with the clash score, indicated that these base pairs adopt a Hoogsteen conformation.

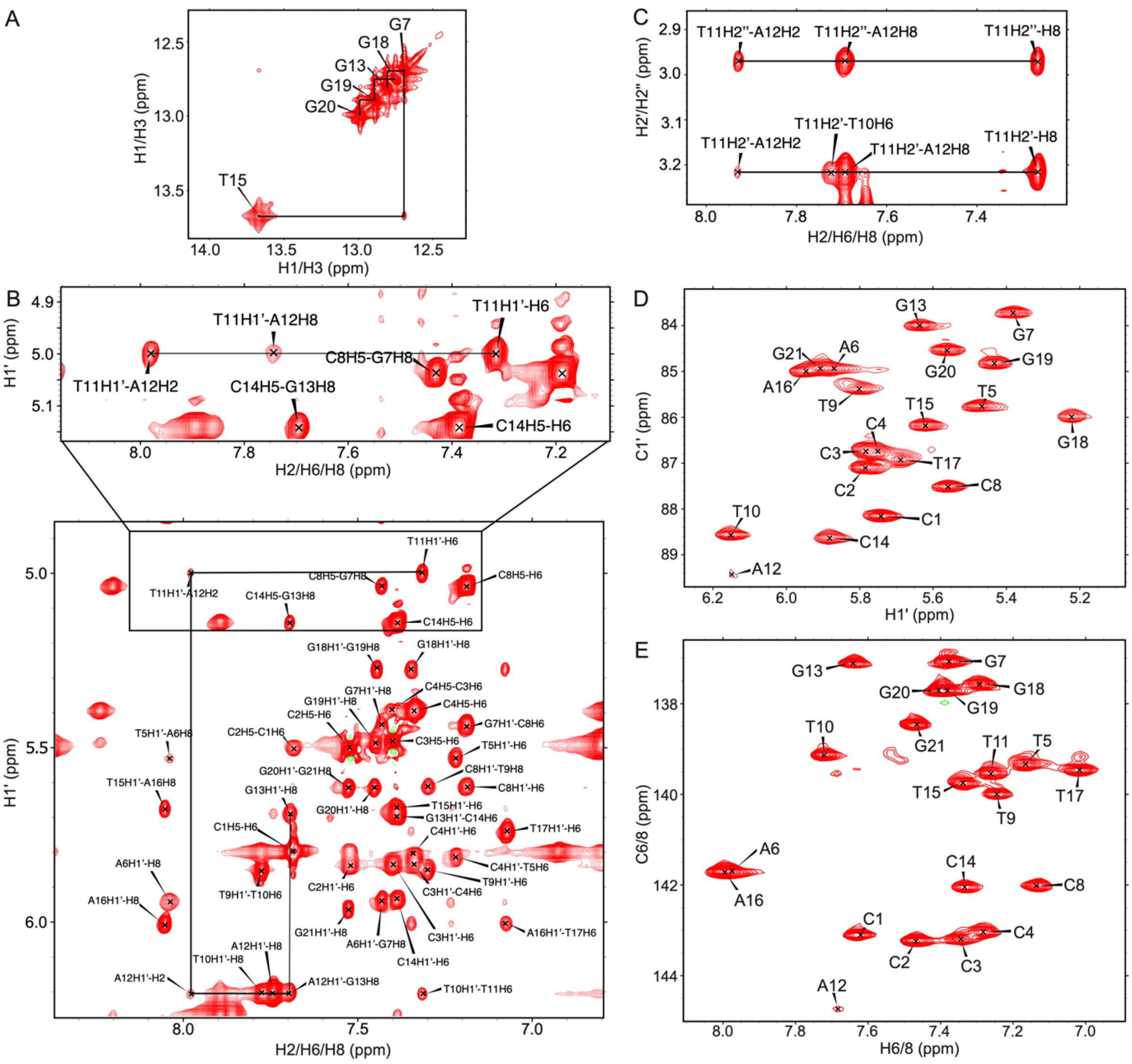


**Figure S3 Resonance assignment of RE construct.** (A-C) Selected regions of the 2D $^1$H-$^1$H NOESY spectrum of RE with mixing time 200 ms. Shown are the (A) H1/H3; (B) H1'-H8/H6/H2; and (C) H2'/H2''-H8/H6/H2 regions. Solid lines indicate the NOE-walk used for sequential assignment. (D-E) 2D $^1$H-$^{13}$C HSQC NMR spectra showing the (D) sugar H1'-C1'; and (E) aromatic H6/H8-C6/C8 regions.

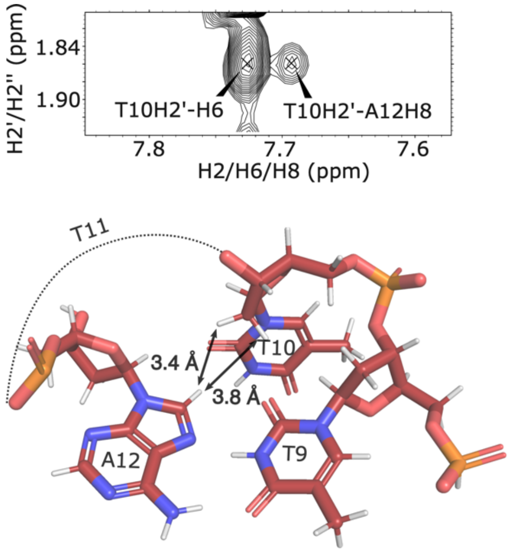


**Figure S4. A long-range NOE contact for RE in solution is consistent with the A12-T9 base pair forming a Hoogsteen conformation.** Selected region of the 2D $^1$H-$^1$H 2D NOESY spectra (top, mixing time 200 ms) of RE showing a long-range A12(H8)···T10(H2') NOE cross-peak, which is consistent with distances derived from the crystal structure (bottom, black arrows), and with A12 adopting the *syn* but not the *anti* conformation.

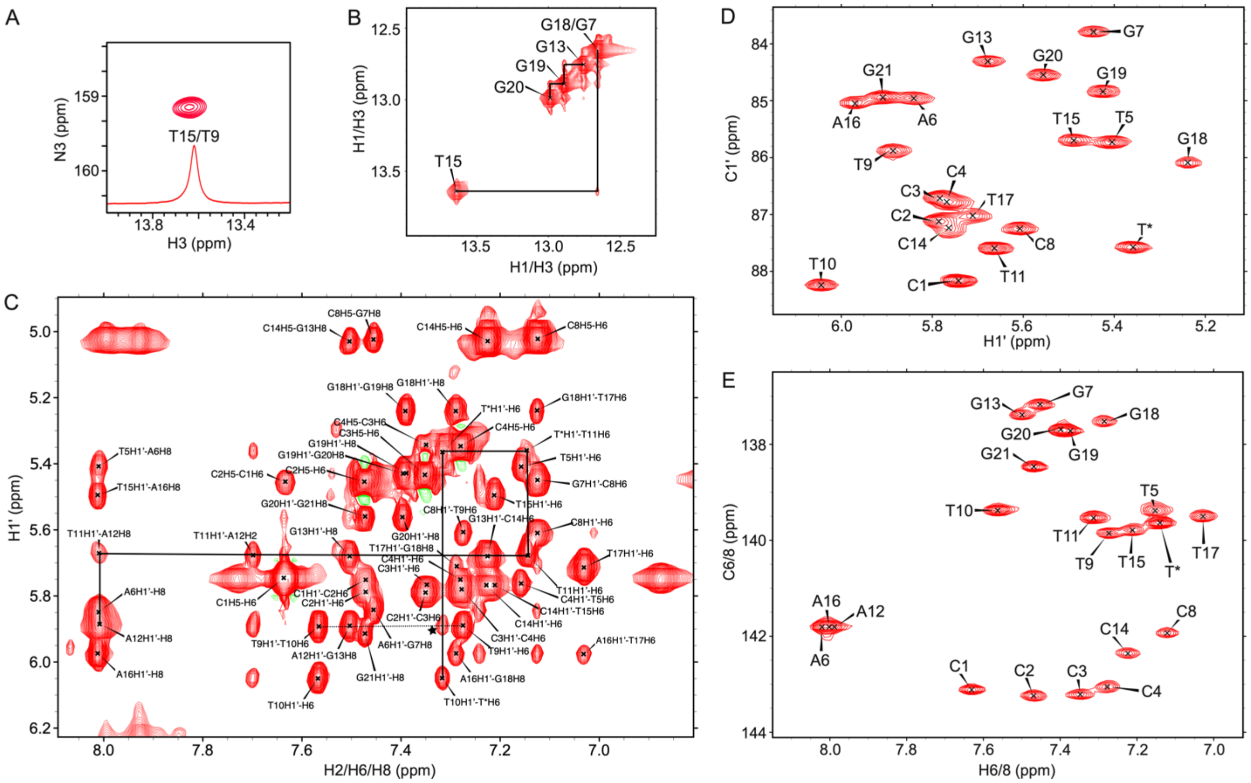

**Figure S5. Resonance assignment of LE construct.** (A) 2D $^{1}$H-$^{15}$N HSQC of site labelled A12-T9 LE shows T9(N3) and T9(H3) chemical shifts typical for a Watson-Crick conformation. Overlayed is the corresponding 1D $^{1}$H imino spectrum. (B-C) Selected regions of the 2D $^{1}$H-$^{1}$H NOESY spectrum of LE with mixing time 200 ms. Shown are the (B) H1/H3 and (C) H1'-H8/H6/H2 regions. Solid lines indicate the NOE-walk used for sequential assignment. (D-E) 2D $^{1}$H-$^{13}$C HSQC NMR spectra showing the (D) sugar H1'-C1'; and (E) aromatic H6/H8-C6/C8 regions.

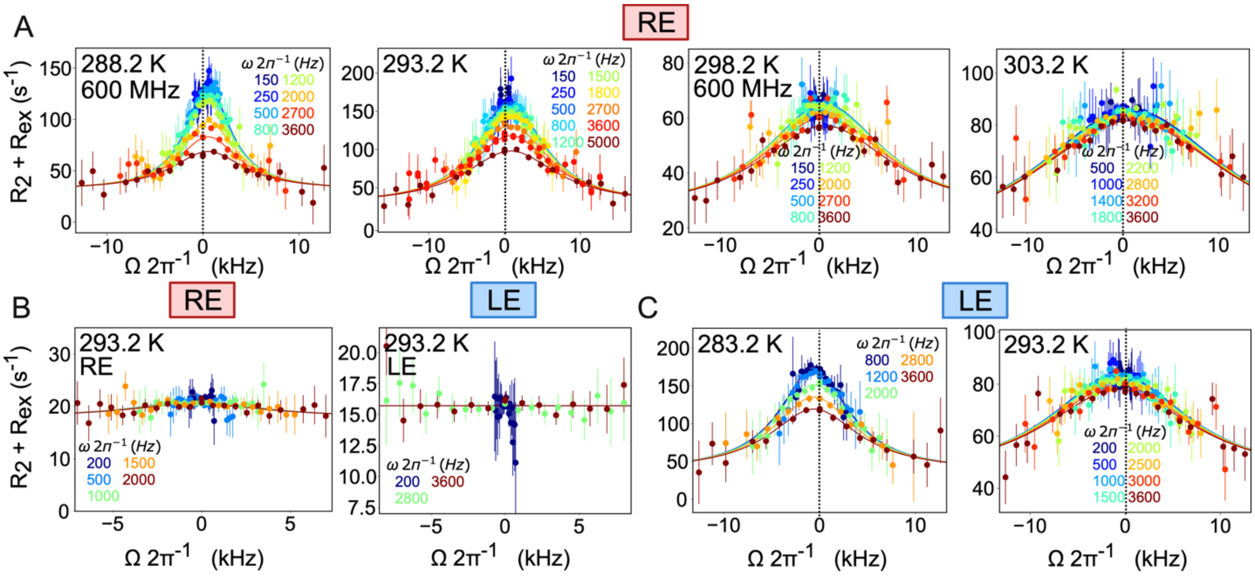


**Figure S6. Additional off-resonance $^{13}$C $R_{1\rho}$ profiles.** Off-resonance $^{13}$C $R_{1\rho}$ profiles measured at different spin-lock powers (color-coded) on (A) A12(C8) for RE; (B) T9(C1') for RE (right) and LE (left); and (C) A12(C8) for LE at 900 MHz, unless otherwise stated. Shown is the fit (solid line) to a 2-state exchange model using Bloch–McConnell equations, as described in Methods. Error bars represent the experimental uncertainty in the $R_{1\rho}$ data and were obtained by propagating the error in $R_{1\rho}$ using the Monte Carlo procedure. Dashed vertical line at 0 Hz offset shows how profiles are displaced along opposite directions for RE and LE, consistent with the inversion of the chemical shifts of the major and minor states.

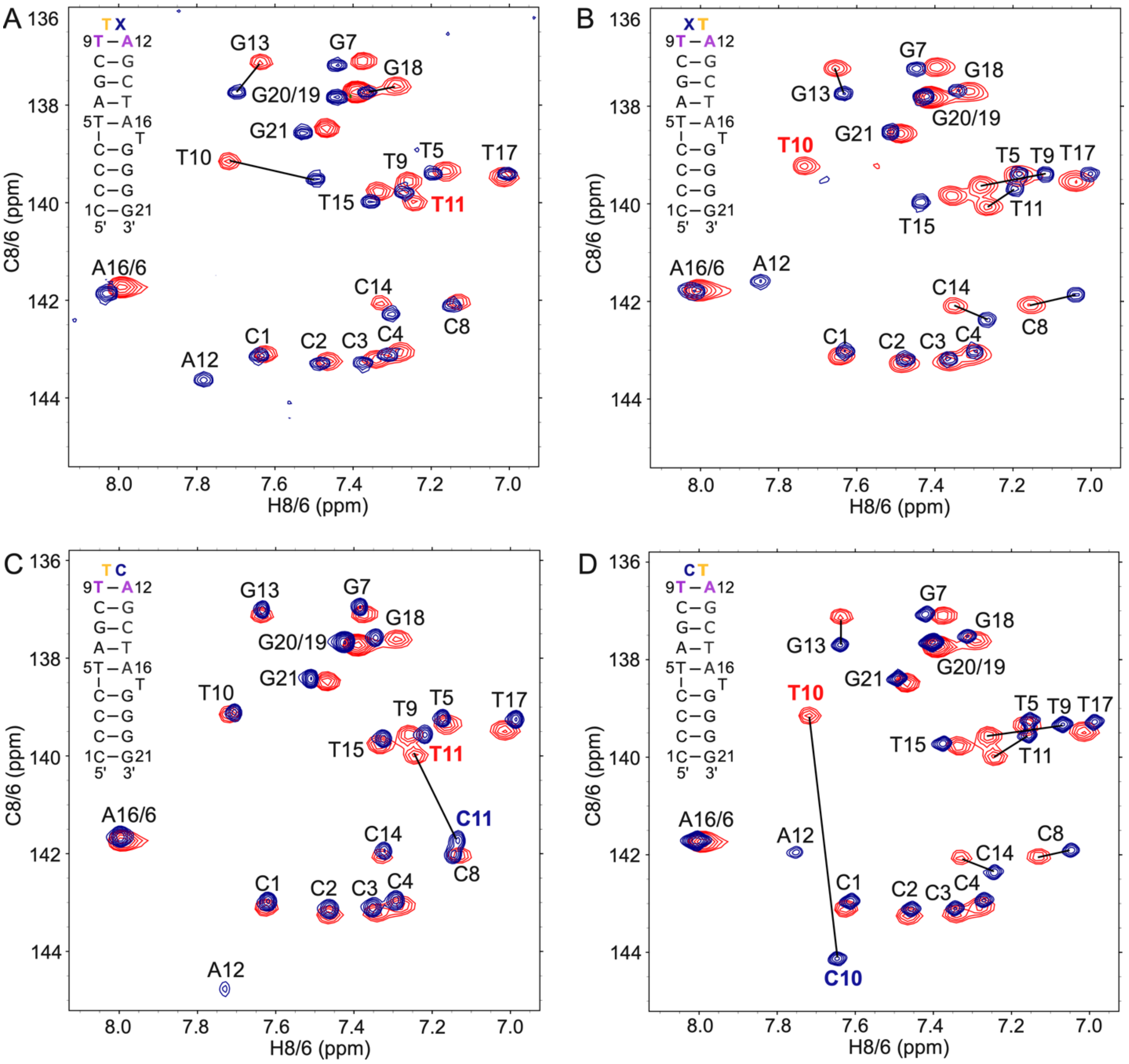


**Figure S7. Substitutions in the apical loop residues shift the A12-T9 Watson-Crick-Hoogsteen equilibrium.** Aromatic (H6/H8-C6/C8) regions from 2D $^{1}H$-$^{13}C$ HSQC spectra of the RE mutants (blue) together with the corresponding DNA construct: (A) Ab11, (B) Ab10, (C) TC, (D) CT, overlayed with RE (red).

| Nucleotide position: | 8 | 9 | 10 | 11 | 12 | 13 |
|---|---|---|---|---|---|---|
| A | 0 | 0 | 0 | 0 | 768 | 0 |
| C | 768 | 0 | 0 | 8 | 0 | 0 |
| G | 0 | 0 | 0 | 0 | 0 | 768 |
| T | 0 | 768 | 768 | 760 | 0 | 0 |

**Figure S8. Position weighted matrix of the RE apical loop from IS608.** IS608 transposon sequences were identified and aligned using BLAST[18] in *Helicobacter pylori* using a reference sequence from IS database (https://isfinder.biotoul.fr)[19]. Only *Helicobacter pylori* sequences with ≥98% coverage were retained. The apical loop region of the RE hairpin (positions #8-13) was then isolated, to count nucleotide frequencies at each position and generate the position weighted matrix.

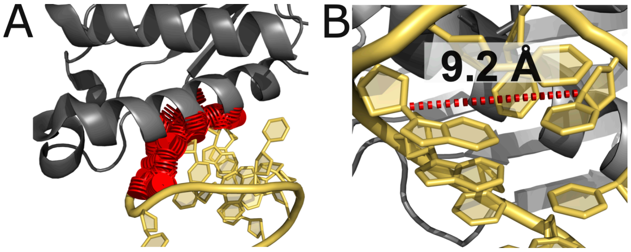


**Figure S9. Fitting of the hairpin with A12-T9 base pair in Watson-Crick conformation results in clashes with TnpA.** (A) Modelling the apical loop with an A12(*anti*)-T9 Watson-Crick conformation resulted in the loss of critical contacts and introduced steric clashes. (B) Representative AlphaFold3 predicted structure prediction for a TnpA-LE complex exhibiting an A12(*anti*)-T9 base pair with constricted C1'–C1' distances <10 Å.

**Table S1.** Oligonucleotides sequences used in this study. In the RE and LE constructs, residues A12 and T9 (in bold) were either unlabeled, or uniformly $^{13}C/^{15}N$-labeled (see Methods). In the modified constructs, RE loop nucleotides T10/T11 were replaced with an abasic site (Ab) or a cytosine (C). The site replaced is highlighted in bold.

| Construct Name | Sequence (5' – 3') |
|---|---|
| RE | CCCCTAGC**T**TT**A**GCTATGGGG |
| LE | CCCCTAGC**T**TTT**A**GCTATGGGG |
| Ab10 | CCCCTAGCT**Ab**TAGCTATGGGG |
| Ab11 | CCCCTAGCTT**Ab**AGCTATGGGG |
| TC | CCCCTAGCTT**C**AGCTATGGGG |
| CT | CCCCTAGCT**C**TAGCTATGGGG |

**Table S2.** NOE contacts in RE and LE and their comparison with corresponding PDB structures (PDB ID: 2VJV, 2VGH). Ambiguous peak assignments are indicated by "/". The expected contact was either observed (+), or not observed (-) by NMR.

| Nucleotide1 | Atom1 | Nucleotide2 | Atom2 | PDB distance, Å | NMR RE | NMR LE |
|---|---|---|---|---|---|---|
| G13 | H8 | A12 | H1' | 4.4 | A12/T10 | + |
| G13 | H8 | A12 | H2' | 3.4 | + | + |
| G13 | H1' | A12 | H8 | 4.6 | + | G13/T* |
| G13 | H1' | A12 | H2 | >10 | - | + |
| T9 | H6 | C8 | H1' | 3 | + | + |
| T9 | H6 | C8 | H2' | 2.2 | + | + |
| T9 | H6 | C8 | H2'' | 3.8 | - | - |
| T9 | H7 | C8 | H1' | 3.6 | + | + |
| A12 | H8 | A12 | H1' | 2.5 | + | + |
| A12 | H8 | T9 | H3 | 3.5 | + | - |
| A12 | H8 | T10 | H2' | 3.8 | - | - |
| A12 | H8 | T10 | H2'' | 3.4 | + | - |
| A12 | H8 | T10 | H1' | 4.2 | A12/T10 | - |
| A12 | H2 | T10 | H1' | 9.5 | A12/T10 | + |
| A12 | H2 | T10 | H2' | 7.8 | - | + |
| A12 | H2 | T10 | H2'' | 8.9 | + | + |
| A12 | H2 | T11 | H2' | 5.9 | + | + |
| A12 | H2 | T11 | H2'' | 6.9 | + | - |
| A12 | H2 | T11 | H1' | 4.8 | + | + |
| A12 | H2 | T11 | H6 | 7.9 | - | - |
| A12 | H8 | T11 | H2' | 6.7 | + | - |
| A12 | H8 | T11 | H2'' | 7.3 | + | - |
| A12 | H8 | T11 | H1' | 5.8 | + | + |
| A12 | H8 | T11 | H6 | 5.5 | - | - |
| A12 | H1' | T11 | H6 | 6.7 | A12/T10 | A12/T9 |
| A12 | H1' | T10 | H6 | 6.9 | A12/T10 | A12/T9 |

| | | | | | | |
|---|---|---|---|---|---|---|
| T11 | H6 | T10 | H2' | 3.4 | + | + |
| T11 | H6 | T10 | H2" | 3.3 | - | + |
| T11 | H6 | T10 | H1' | 5.6 | A12/T10 | + |
| T11 | H7 | T10 | H1' | 6 | A12/T10 | - |
| T11 | H7 | T9 | H1' | 4.1 | - | A12/T9 |
| T10 | H6 | T9 | H2' | 4.1 | + | + |
| T10 | H6 | T9 | H2" | 5.4 | - | - |
| T10 | H6 | T9 | H1' | 3.2 | + | A12/T9 |
| T10 | H6 | T11 | H2' | 8.9 | + | + |
| T10 | H6 | G13 | H1' | 3.1 | - | - |
| T10 | H6 | G13 | H8 | 5.5 | - | - |
| T10 | H7 | T10 | H1' | 5.7 | A12/T10 | + |
| T10 | H7 | T9 | H1' | 3.3 | + | A12/T9 |
| T10 | H7 | C8 | H6 | >10 | + | - |
| T10 | H7 | C8 | H1' | 5.7 | - | + |
| T10 | H7 | C14 | H1' | 6.6 | - | + |
| T10 | H7 | A12 | H2 | >10 | - | + |
| T* | H6 | T11 | H1' | >10 | N/A | + |
| T* | H1' | A12 | H2 | 6.4 | N/A | G13/T* |
| T* | H1' | A12 | H8 | >10 | N/A | + |
| T* | H2' | A12 | H2 | 5.9 | N/A | + |
| T* | H2" | A12 | H2 | 6.5 | N/A | + |
| T* | H2" | A12 | H8 | 9.7 | N/A | + |

**Table S3.** List of spin-lock powers ($\omega_1/2\pi$ in Hz) and offsets ($\Omega_{eff}/2\pi$ in Hz) used in off-resonance $^{13}$C $R_{1\rho}$ experiments of RE.

| Residue | Conditions | $\omega_1/2\pi$ (Hz), [$\Omega_{eff}/2\pi$ (Hz)] |
|---|---|---|
| **RE A12(C8)** | 15 °C 900 MHz | 200, [-704.0, -640.0, -576.0, -512.0, -448.0, -384.0, -320.0, -256.0, -192.0, -128.0, -64.0, -10.0, 10.0, 64.0, 128.0, 192.0, 256.0, 320.0, 384.0, 448.0, 512.0, 576.0, 640.0, 704.0] |
| | | 800, [-2805.0, -2550.0, -2295.0, -2040.0, -1785.0, -1530.0, -1275.0, -1020.0, -765.0, -510.0, -255.0, -10.0, 10.0, 255.0, 510.0, 765.0, 1020.0, 1275.0, 1530.0, 1785.0, 2040.0, 2295.0, 2550.0, 2805.0] |
| | | 1500, [-5247.0, -4770.0, -4293.0, -3816.0, -3339.0, -2862.0, -2385.0, -1908.0, -1431.0, -954.0, -477.0, -10.0, 10.0, 477.0, 954.0, 1431.0, 1908.0, 2385.0, 2862.0, 3339.0, 3816.0, 4293.0, 4770.0, 5247.0] |
| | | 2400, [-8404.0, -7640.0, -6876.0, -6112.0, -5348.0, -4584.0, -3820.0, -3056.0, -2292.0, -1528.0, -764.0, -10.0, 10.0, 764.0, 1528.0, 2292.0, 3056.0, 3820.0, 4584.0, 5348.0, 6112.0, 6876.0, 7640.0, 8404.0] |
| | | 3000, [-10505.0, -9550.0, -8595.0, -7640.0, -6685.0, -5730.0, -4775.0, -3820.0, -2865.0, -1910.0, -955.0, -10.0, 10.0, 955.0, 1910.0, 2865.0, 3820.0, 4775.0, 5730.0, 6685.0, 7640.0, 8595.0, 9550.0, 10505.0] |
| | | 3600, [-12595.0, -11450.0, -10305.0, -9160.0, -8015.0, -6870.0, -5725.0, -4580.0, -3435.0, -2290.0, -1145.0, -10.0, 10.0, 1145.0, 2290.0, 3435.0, 4580.0, 5725.0, 6870.0, 8015.0, 9160.0, 10305.0, 11450.0, 12595.0] |
| | 20 °C 900 MHz | 150, [-528.0, -480.0, -432.0, -384.0, -336.0, -288.0, -240.0, -192.0, -144.0, -96.0, -48.0, -10.0, 10.0, 48.0, 96.0, 144.0, 192.0, 240.0, 288.0, 336.0, 384.0, 432.0, 480.0, 528.0] |

| | | |
|---|---|---|
| | | 250, [-880.0, -800.0, -720.0, -640.0, -560.0, -480.0, -400.0, -320.0, -240.0, -160.0, -80.0, -10.0, 10.0, 80.0, 160.0, 240.0, 320.0, 400.0, 480.0, 560.0, 640.0, 720.0, 800.0, 880.0]<br>500, [-1749.0, -1590.0, -1431.0, -1272.0, -1113.0, -954.0, -795.0, -636.0, -477.0, -318.0, -159.0, -10.0, 10.0, 159.0, 318.0, 477.0, 636.0, 795.0, 954.0, 1113.0, 1272.0, 1431.0, 1590.0, 1749.0]<br>800, [-2805.0, -2805.0, -2550.0, -2550.0, -2295.0, -2295.0, -2040.0, -2040.0, -1785.0, -1785.0, -1530.0, -1530.0, -1275.0, -1275.0, -1020.0, -1020.0, -765.0, -765.0, -510.0, -510.0, -255.0, -255.0, -10.0, -10.0, 10.0, 10.0, 255.0, 255.0, 510.0, 510.0, 765.0, 765.0, 1020.0, 1020.0, 1275.0, 1275.0, 1530.0, 1530.0, 1785.0, 1785.0, 2040.0, 2040.0, 2295.0, 2295.0, 2550.0, 2550.0, 2805.0, 2805.0]<br>1200, [-4202.0, -3820.0, -3438.0, -3056.0, -2674.0, -2292.0, -1910.0, -1528.0, -1146.0, -764.0, -382.0, -10.0, 10.0, 382.0, 764.0, 1146.0, 1528.0, 1910.0, 2292.0, 2674.0, 3056.0, 3438.0, 3820.0, 4202.0]<br>1500, [-5247.0, -4770.0, -4293.0, -3816.0, -3339.0, -2862.0, -2385.0, -1908.0, -1431.0, -954.0, -477.0, -10.0, 10.0, 477.0, 954.0, 1431.0, 1908.0, 2385.0, 2862.0, 3339.0, 3816.0, 4293.0, 4770.0, 5247.0]<br>1800, [-6303.0, -5730.0, -5157.0, -4584.0, -4011.0, -3438.0, -2865.0, -2292.0, -1719.0, -1146.0, -573.0, -10.0, 10.0, 573.0, 1146.0, 1719.0, 2292.0, 2865.0, 3438.0, 4011.0, 4584.0, 5157.0, 5730.0, 6303.0]<br>2700, [-9449.0, -7731.0, -6872.0, -6013.0, -5154.0, -4295.0, -3436.0, -2577.0, -1718.0, -859.0, -10.0, 10.0, 859.0, 1718.0, 2577.0, 3436.0, 4295.0, 5154.0, 6013.0, 6872.0, 7731.0]<br>3600, [-12595.0, -12595.0, -11450.0, -11450.0, -10305.0, -10305.0, -9160.0, -8015.0, -6870.0, -6870.0, -5725.0, -5725.0, -4580.0, -4580.0, -3435.0, -3435.0, - |

| | | |
|---|---|---|
| | | 2290.0, -2290.0, -1145.0, -1145.0, -10.0, -10.0, 10.0, 10.0, 1145.0, 1145.0, 2290.0, 2290.0, 3435.0, 3435.0, 4580.0, 4580.0, 5725.0, 5725.0, 6870.0, 6870.0, 8015.0, 9160.0, 10305.0, 10305.0, 11450.0, 11450.0, 12595.0, 12595.0]<br>5000, [-15910.0, -14319.0, -12728.0, -11137.0, -9546.0, -6364.0, -4773.0, -3182.0, -1591.0, -10.0, 10.0, 1591.0, 3182.0, 4773.0, 6364.0, 9546.0, 11137.0, 12728.0, 14319.0, 15910.0] |
| | 30 °C<br>900 MHz | 500, [-1749.0, -1590.0, -1272.0, -1113.0, -954.0, -795.0, -636.0, -477.0, -318.0, -159.0, -10.0, 10.0, 159.0, 318.0, 477.0, 636.0, 795.0, 954.0, 1113.0, 1272.0, 1431.0, 1590.0, 1749.0]<br>1000, [-3498.0, -3180.0, -2862.0, -2544.0, -2226.0, -1908.0, -1590.0, -1272.0, -954.0, -636.0, -318.0, -10.0, 10.0, 318.0, 636.0, 954.0, 1272.0, 1590.0, 1908.0, 2226.0, 2544.0, 2862.0, 3180.0, 3498.0]<br>1400, [-4895.0, -4450.0, -4005.0, -3560.0, -3115.0, -2670.0, -2225.0, -1780.0, -1335.0, -890.0, -445.0, -10.0, 10.0, 445.0, 890.0, 1335.0, 1780.0, 2225.0, 2670.0, 3115.0, 3560.0, 4005.0, 4450.0, 4895.0]<br>1800, [-6303.0, -5730.0, -5157.0, -4584.0, -4011.0, -3438.0, -2865.0, -2292.0, -1719.0, -1146.0, -573.0, -10.0, 10.0, 573.0, 1146.0, 1719.0, 2292.0, 2865.0, 3438.0, 4011.0, 4584.0, 5157.0, 5730.0, 6303.0]<br>2200, [-7700.0, -7000.0, -6300.0, -5600.0, -4900.0, -4200.0, -3500.0, -2800.0, -2100.0, -1400.0, -700.0, -10.0, 10.0, 700.0, 1400.0, 2100.0, 2800.0, 3500.0, 4200.0, 4900.0, 5600.0, 6300.0, 7000.0, 7700.0]<br>2800, [-9801.0, -8019.0, -7128.0, -6237.0, -5346.0, -4455.0, -3564.0, -2673.0, -1782.0, -891.0, -10.0, 10.0, 891.0, 1782.0, 2673.0, 3564.0, 4455.0, 5346.0, 6237.0, 7128.0, 8019.0, 9801.0] |

| | | |
|---|---|---|
| | | 3200, [-11198.0, -10180.0, -7126.0, -6108.0, -5090.0, -4072.0, -3054.0, -2036.0, -1018.0, -10.0, 10.0, 1018.0, 2036.0, 3054.0, 4072.0, 5090.0, 6108.0, 7126.0, 10180.0, 11198.0]<br>3600, [-12595.0, -11450.0, -10305.0, -6870.0, -5725.0, -4580.0, -3435.0, -2290.0, -1145.0, -10.0, 10.0, 1145.0, 2290.0, 3435.0, 4580.0, 5725.0, 6870.0, 10305.0, 11450.0, 12595.0] |
| | 15 °C<br>600 MHz | 150, [-528.0, -480.0, -432.0, -384.0, -336.0, -288.0, -240.0, -192.0, -144.0, -96.0, -48.0, -10.0, 10.0, 48.0, 96.0, 144.0, 192.0, 240.0, 288.0, 336.0, 384.0, 432.0, 480.0, 528.0]<br>250, [-880.0, -800.0, -720.0, -640.0, -560.0, -480.0, -400.0, -320.0, -240.0, -160.0, -80.0, -10.0, 10.0, 80.0, 160.0, 240.0, 320.0, 400.0, 480.0, 560.0, 640.0, 720.0, 800.0, 880.0]<br>500, [-1749.0, -1590.0, -1431.0, -1272.0, -1113.0, -954.0, -795.0, -636.0, -477.0, -318.0, -159.0, -10.0, 10.0, 159.0, 318.0, 477.0, 636.0, 795.0, 954.0, 1113.0, 1272.0, 1431.0, 1590.0, 1749.0]<br>800, [-2805.0, -2550.0, -2295.0, -2040.0, -1785.0, -1530.0, -1275.0, -1020.0, -765.0, -510.0, -255.0, -10.0, 10.0, 255.0, 510.0, 765.0, 1020.0, 1275.0, 1530.0, 1785.0, 2040.0, 2295.0, 2550.0, 2805.0]<br>1200, [-4202.0, -3820.0, -3438.0, -3056.0, -2674.0, -2292.0, -1910.0, -1528.0, -1146.0, -764.0, -382.0, -10.0, 10.0, 382.0, 764.0, 1146.0, 1528.0, 1910.0, 2292.0, 2674.0, 3056.0, 3438.0, 3820.0, 4202.0]<br>2000, [-6996.0, -6360.0, -5724.0, -5088.0, -4452.0, -3816.0, -3180.0, -2544.0, -1908.0, -1272.0, -636.0, -10.0, 10.0, 636.0, 1272.0, 1908.0, 2544.0, 3180.0, 3816.0, 4452.0, 5088.0, 5724.0, 6360.0, 6996.0] |

| | | |
|---|---|---|
| | | 2700, [-8590.0, -7731.0, -6872.0, -6013.0, -5154.0, -4295.0, -3436.0, -2577.0, -1718.0, -859.0, -10.0, 10.0, 859.0, 1718.0, 2577.0, 3436.0, 4295.0, 5154.0, 6013.0, 6872.0, 7731.0, 8590.0]<br>3600, [-12595.0, -11450.0, -10305.0, -8015.0, -6870.0, -5725.0, -4580.0, -3435.0, -2290.0, -1145.0, -10.0, 10.0, 1145.0, 2290.0, 3435.0, 4580.0, 5725.0, 6870.0, 8015.0, 10305.0, 11450.0, 12595.0] |
| | 25 °C<br>600 MHz | [150], {-528.0, -480.0, -432.0, -384.0, -336.0, -288.0, -240.0, -192.0, -144.0, -96.0, -48.0, -10.0, 10.0, 48.0, 96.0, 144.0, 192.0, 240.0, 288.0, 336.0, 384.0, 432.0, 480.0, 528.0}<br>[250], {-880.0, -800.0, -720.0, -640.0, -560.0, -480.0, -400.0, -320.0, -240.0, -160.0, -80.0, -10.0, 10.0, 80.0, 160.0, 240.0, 320.0, 400.0, 480.0, 560.0, 640.0, 720.0, 800.0, 880.0}<br>[500], {-1749.0, -1590.0, -1431.0, -1272.0, -1113.0, -954.0, -795.0, -636.0, -477.0, -318.0, -159.0, -10.0, 10.0, 159.0, 318.0, 477.0, 636.0, 795.0, 954.0, 1113.0, 1272.0, 1431.0, 1590.0, 1749.0}<br>[800], {-2805.0, -2550.0, -2295.0, -2040.0, -1785.0, -1530.0, -1275.0, -1020.0, -765.0, -510.0, -255.0, -10.0, 10.0, 255.0, 510.0, 765.0, 1020.0, 1275.0, 1530.0, 1785.0, 2040.0, 2295.0, 2550.0, 2805.0}<br>[1200], {-4202.0, -3820.0, -3438.0, -3056.0, -2674.0, -2292.0, -1910.0, -1528.0, -1146.0, -764.0, -382.0, -10.0, 10.0, 382.0, 764.0, 1146.0, 1528.0, 1910.0, 2292.0, 2674.0, 3056.0, 3438.0, 3820.0, 4202.0}<br>[2000], {-6996.0, -6360.0, -5724.0, -5088.0, -4452.0, -3816.0, -3180.0, -2544.0, -1908.0, -1272.0, -636.0, -10.0, 10.0, 636.0, 1272.0, 1908.0, 2544.0, 3180.0, 3816.0, 4452.0, 5088.0, 5724.0, 6360.0, 6996.0} |

| | | |
|---|---|---|
| | | [2700], {-8590.0, -7731.0, -6872.0, -6013.0, -5154.0, -4295.0, -3436.0, -2577.0, -1718.0, -859.0, -10.0, 10.0, 859.0, 1718.0, 2577.0, 3436.0, 4295.0, 5154.0, 6013.0, 6872.0, 7731.0, 8590.0}<br>[3600], {-12595.0, -11450.0, -10305.0, -8015.0, -6870.0, -5725.0, -4580.0, -3435.0, -2290.0, -1145.0, -10.0, 10.0, 1145.0, 2290.0, 3435.0, 4580.0, 5725.0, 6870.0, 8015.0, 10305.0, 11450.0, 12595.0} |
| **RE A12(C1')** | 15 °C<br>600 MHz | 150, [-528.0, -480.0, -432.0, -384.0, -336.0, -288.0, -240.0, -192.0, -144.0, -96.0, -48.0, -10.0, 10.0, 48.0, 96.0, 144.0, 192.0, 240.0, 288.0, 336.0, 384.0, 432.0, 480.0, 528.0]<br>800, [-2805.0, -2550.0, -2295.0, -2040.0, -1785.0, -1530.0, -1275.0, -1020.0, -765.0, -510.0, -255.0, -10.0, 10.0, 255.0, 510.0, 765.0, 1020.0, 1275.0, 1530.0, 1785.0, 2040.0, 2295.0, 2550.0, 2805.0]<br>1200, [-4202.0, -3820.0, -3438.0, -3056.0, -2674.0, -2292.0, -1910.0, -1528.0, -1146.0, -764.0, -382.0, -10.0, 10.0, 382.0, 764.0, 1146.0, 1528.0, 1910.0, 2292.0, 2674.0, 3056.0, 3438.0, 3820.0, 4202.0]<br>1600, [-5599.0, -5090.0, -4581.0, -4072.0, -3563.0, -3054.0, -2545.0, -2036.0, -1527.0, -1018.0, -509.0, -10.0, 10.0, 509.0, 1018.0, 1527.0, 2036.0, 2545.0, 3054.0, 3563.0, 4072.0, 4581.0, 5090.0, 5599.0]<br>2000, [-6996.0, -6360.0, -5724.0, -5088.0, -4452.0, -3816.0, -3180.0, -2544.0, -1908.0, -1272.0, -636.0, -10.0, 10.0, 636.0, 1272.0, 1908.0, 2544.0, 3180.0, 3816.0, 4452.0, 5088.0, 5724.0, 6360.0, 6996.0]<br>2400, [-7640.0, -6876.0, -6112.0, -5348.0, -4584.0, -3820.0, -3056.0, -2292.0, -1528.0, -764.0, -10.0, 10.0, 764.0, 1528.0, 2292.0, 3056.0, 3820.0, 4584.0, 5348.0, 6112.0, 6876.0, 7640.0, 8404.0] |

| | | |
|---|---|---|
| | | 2800, [-8910.0, -7128.0, -6237.0, -5346.0, -4455.0, -3564.0, -2673.0, -1782.0, -891.0, -10.0, 10.0, 891.0, 1782.0, 2673.0, 3564.0, 4455.0, 5346.0, 6237.0, 7128.0, 8910.0, 9801.0]<br>3600, [-12595.0, -11450.0, -10305.0, -8015.0, -6870.0, -5725.0, -4580.0, -3435.0, -2290.0, -1145.0, -10.0, 10.0, 1145.0, 2290.0, 3435.0, 4580.0, 5725.0, 6870.0, 10305.0, 11450.0, 12595.0] |
| **RE A12(C2)** | 20 °C 900 MHz | 200, [-704.0, -640.0, -576.0, -512.0, -448.0, -384.0, -320.0, -256.0, -192.0, -128.0, -64.0, -10.0, 10.0, 64.0, 128.0, 192.0, 256.0, 320.0, 384.0, 448.0, 512.0, 576.0, 640.0, 704.0]<br>500, [-1749.0, -1590.0, -1431.0, -1272.0, -1113.0, -954.0, -795.0, -636.0, -477.0, -318.0, -159.0, -10.0, 10.0, 159.0, 318.0, 477.0, 636.0, 795.0, 954.0, 1113.0, 1272.0, 1431.0, 1590.0, 1749.0]<br>1000, [-3498.0, -3180.0, -2862.0, -2544.0, -2226.0, -1908.0, -1590.0, -1272.0, -954.0, -636.0, -318.0, -10.0, 10.0, 318.0, 636.0, 954.0, 1272.0, 1590.0, 1908.0, 2226.0, 2544.0, 2862.0, 3180.0, 3498.0]<br>1500, [-5247.0, -4770.0, -4293.0, -3816.0, -3339.0, -2862.0, -2385.0, -1908.0, -1431.0, -954.0, -477.0, -10.0, 10.0, 477.0, 954.0, 1431.0, 1908.0, 2385.0, 2862.0, 3339.0, 3816.0, 4293.0, 4770.0, 5247.0]<br>2000, [-6996.0, -6360.0, -5724.0, -5088.0, -4452.0, -3816.0, -3180.0, -2544.0, -1908.0, -1272.0, -636.0, -10.0, 10.0, 636.0, 1272.0, 1908.0, 2544.0, 3180.0, 3816.0, 4452.0, 5088.0, 5724.0, 6360.0, 6996.0] |
| **RE T9(C6)** | 20 °C 900 MHz | 200, [-704.0, -640.0, -576.0, -512.0, -448.0, -384.0, -320.0, -256.0, -192.0, -128.0, -64.0, -10.0, 10.0, 64.0, 128.0, 192.0, 256.0, 320.0, 384.0, 448.0, 512.0, 576.0, 640.0, 704.0] |

| | | |
|---|---|---|
| | | 500, [-1749.0, -1590.0, -1431.0, -1272.0, -1113.0, -954.0, -795.0, -636.0, -477.0, -318.0, -159.0, -10.0, 10.0, 159.0, 318.0, 477.0, 636.0, 795.0, 954.0, 1113.0, 1272.0, 1431.0, 1590.0, 1749.0]<br>1000, [-3498.0, -2862.0, -2544.0, -2226.0, -1908.0, -1590.0, -1272.0, -954.0, -636.0, -318.0, -10.0, 10.0, 318.0, 636.0, 954.0, 1272.0, 1590.0, 1908.0, 2226.0, 2544.0, 2862.0, 3180.0, 3498.0]<br>1500, [-5247.0, -4770.0, -4293.0, -3816.0, -3339.0, -2862.0, -2385.0, -1908.0, -1431.0, -954.0, -477.0, -10.0, 10.0, 477.0, 954.0, 1431.0, 1908.0, 2385.0, 2862.0, 3339.0, 3816.0, 4293.0, 4770.0, 5247.0]<br>2000, [-6996.0, -6360.0, -5724.0, -5088.0, -4452.0, -3816.0, -2544.0, -1908.0, -1272.0, -636.0, -10.0, 10.0, 636.0, 1272.0, 1908.0, 2544.0, 3180.0, 3816.0, 4452.0, 5088.0, 5724.0, 6360.0, 6996.0]<br>3000, [-10505.0, -9550.0, -8595.0, -7640.0, -6685.0, -5730.0, -4775.0, -3820.0, -2865.0, -1910.0, -955.0, -10.0, 10.0, 955.0, 1910.0, 2865.0, 3820.0, 4775.0, 5730.0, 6685.0, 7640.0, 8595.0, 9550.0, 10505.0] |
| **RE T9(C1')** | 20 °C<br>900 MHz | 200, [-704.0, -640.0, -576.0, -512.0, -448.0, -384.0, -320.0, -256.0, -192.0, -128.0, -64.0, -10.0, 10.0, 64.0, 128.0, 192.0, 256.0, 320.0, 384.0, 448.0, 512.0, 576.0, 640.0, 704.0]<br>500, [-1749.0, -1590.0, -1431.0, -1272.0, -1113.0, -954.0, -795.0, -636.0, -477.0, -318.0, -159.0, -10.0, 10.0, 159.0, 318.0, 477.0, 636.0, 795.0, 954.0, 1113.0, 1272.0, 1431.0, 1590.0, 1749.0]<br>1000, [-3498.0, -3180.0, -2862.0, -2544.0, -2226.0, -1908.0, -1590.0, -1272.0, -954.0, -636.0, -318.0, -10.0, 10.0, 318.0, 636.0, 954.0, 1272.0, 1590.0, 1908.0, 2226.0, 2544.0, 2862.0, 3180.0, 3498.0] |

| | | |
|---|---|---|
| | | 1500, [-5247.0, -4770.0, -4293.0, -3816.0, -3339.0, -2862.0, -2385.0, -1908.0, -1431.0, -954.0, -477.0, -10.0, 10.0, 477.0, 954.0, 1431.0, 1908.0, 2385.0, 2862.0, 3339.0, 3816.0, 4293.0, 4770.0, 5247.0]<br>2000, [-6996.0, -6360.0, -5724.0, -4452.0, -3816.0, -3180.0, -2544.0, -1908.0, -1272.0, -636.0, -10.0, 10.0, 636.0, 1272.0, 1908.0, 2544.0, 3180.0, 3816.0, 4452.0, 5088.0, 5724.0, 6360.0, 6996.0] |

**Table S4.** Exchange parameters for RE, obtained from global or individual fitting the off-resonance $^{13}$C $R_{1\rho}$ data to the Bloch-McConnell equation. "—" indicates not measured. Error bars represent experimental uncertainty (one s.d.) calculated by the Monte-Carlo approach from a single RD measurement (see Methods).

| Residue | Parameter | 15 °C, 600 MHz | 15 °C, 900 MHz | 20 °C, 900 MHz | 25 °C, 600 MHz | 30 °C, 900 MHz |
|---|---|---|---|---|---|---|
| | | Global Fit | | | | |
| A12 (C8/C1') | $p_{minor,C8}$ (%) | 10.0 ± 0.4 | | 11.5 ± 0.8 | 11 ± 2 | 11 ± 4 |
| | $p_{minor,C1'}$ (%) | | | — | — | — |
| | $k_{ex,C8}$ ($s^{-1}$) | 20500 ± 400 | | 34000 ± 1000 | 46000 ± 3000 | 80000 ± 10000 |
| | $k_{ex,C1'}$ ($s^{-1}$) | | | — | — | — |
| | $R_{1,C8}$ (Hz) | 3.1 ± 0.2 | 2.3± 0.3 | 1.7 ± 0.2 | 2.6 ± 0.1 | 2.2 ± 0.1 |
| | $R_{2,C8}$ (Hz) | 32 ± 1 | 41± 3 | 33 ± 4 | 25 ± 4 | 30 ± 10 |
| | $R_{1,C1'}$ (Hz) | 2.1 ± 0.2 | — | — | — | — |
| | $R_{2,C1'}$ (Hz) | 25 ± 2 | — | — | — | — |
| | $\Delta\omega_{C8}$ (ppm) | -4.7 ± 0.1 | | | | |
| | $\Delta\omega_{C1'}$ (ppm) | 4.6 ± 0.1 | | | | |

| Residue | Parameter | 15 °C, 600 MHz | 15 °C, 900 MHz | 20 °C, 900 MHz | 25 °C, 600 MHz | 30 °C, 900 MHz |
|---|---|---|---|---|---|---|
| | | Individual Fit | | | | |
| A12 (C8/C1') | $p_{minor,C8}$ (%) | 9.9 ± 0.9 | 9.2 ± 0.8 | 12.1 ± 0.9 | 12 ± 3 | 12 ± 5 |
| | $p_{minor,C1'}$ (%) | 10.1 ± 0.6 | — | — | — | — |
| | $k_{ex,C8}$ ($s^{-1}$) | 17800 ± 500 | 20000 ± 700 | 34000 ± 800 | 46000 ± 4000 | 70000 ± 10000 |
| | $k_{ex,C1'}$ ($s^{-1}$) | 17000 ± 500 | — | — | — | — |
| | $R_{1,C8}$ (Hz) | 3.1 ± 0.2 | 2.1 ± 0.4 | 1.7 ± 0.3 | 2.6 ± 0.1 | 2.6 ± 0.4 |
| | $R_{2,C8}$ (Hz) | 30 ± 2 | 44 ± 4 | 34 ± 3 | 25 ± 5 | 38 ± 10 |
| | $R_{1,C1'}$ (Hz) | 2.1 ± 0.2 | — | — | — | — |
| | $R_{2,C1'}$ (Hz) | 24 ± 3 | — | — | — | — |
| | $\Delta\omega_{C8}$ (ppm) | -3.1 ± 0.2 | -4.8 ± 0.1 | -4.5 ± 0.1 | -4.5 ± 0.2 | -4.0 ± 0.6 |
| | $\Delta\omega_{C1'}$ (ppm) | -4.7 ± 0.2 | — | — | — | — |

**Table S5.** List of spin-lock powers ($\omega_1/2\pi$ in Hz) and offsets ($\Omega_{eff}/2\pi$ in Hz) used in off-resonance $^{13}C$ $R_{1\rho}$ experiments of LE.

| **Residue** | **Conditions** | **$\omega_1/2\pi$ (Hz), [$\Omega_{eff}/2\pi$ (Hz)]** |
|---|---|---|
| **LE A12(C8)** | 7.5 °C 900 MHz | 150, [-530.0, -477.0, -424.0, -371.0, -318.0, -265.0, -212.0, -159.0, -106.0, -53.0, -10.0, 10.0, 53.0, 106.0, 159.0, 212.0, 265.0, 318.0, 371.0, 424.0, 477.0, 530.0] |
| | | 500, [-1750.0, -1575.0, -1400.0, -1225.0, -1050.0, -875.0, -700.0, -525.0, -350.0, -175.0, -10.0, 10.0, 175.0, 350.0, 525.0, 700.0, 875.0, 1050.0, 1225.0, 1400.0, 1575.0, 1750.0] |
| | | 1000, [-3500.0, -3150.0, -2800.0, -2450.0, -2100.0, -1750.0, -1400.0, -1050.0, -700.0, -350.0, -10.0, 10.0, 350.0, 700.0, 1050.0, 1400.0, 1750.0, 2100.0, 2450.0, 2800.0, 3150.0, 3500.0] |
| | | 2000, [-7000.0, -6300.0, -5600.0, -4900.0, -4200.0, -3500.0, -2800.0, -2100.0, -1400.0, -700.0, -10.0, 10.0, 700.0, 1400.0, 2100.0, 2800.0, 3500.0, 4200.0, 4900.0, 5600.0, 6300.0, 7000.0] |
| | | 2800, [-9800.0, -7840.0, -6860.0, -5880.0, -4900.0, -3920.0, -2940.0, -1960.0, -980.0, -10.0, 10.0, 980.0, 1960.0, 2940.0, 3920.0, 4900.0, 5880.0, 6860.0, 7840.0, 9800.0] |
| | | 3600, [-12600.0, -11340.0, -10080.0, -7560.0, -6300.0, -5040.0, -3780.0, -2520.0, -1260.0, -10.0, 10.0, 1260.0, 2520.0, 3780.0, 5040.0, 6300.0, 7560.0, 10080.0, 11340.0, 12600.0] |
| | 10 °C 900 MHz | 800, [-2799.0, -2488.0, -2177.0, -1866.0, -1555.0, -1244.0, -933.0, -622.0, -311.0, -10.0, 10.0, 311.0, 622.0, 933.0, 1244.0, 1555.0, 1866.0, 2177.0, 2488.0, 2799.0] |
| | | 1200, [-4203.0, -3736.0, -3269.0, -2802.0, -2335.0, -1868.0, -1401.0, -934.0, -467.0, -10.0, 10.0, 467.0, 934.0, 1401.0, 1868.0, 2335.0, 2802.0, 3269.0, 3736.0, 4203.0] |

| | | |
|---|---|---|
| | | 2000, [-7002.0, -6224.0, -5446.0, -4668.0, -3890.0, -3112.0, -2334.0, -1556.0, -778.0, -10.0, 10.0, 778.0, 1556.0, 2334.0, 3112.0, 3890.0, 4668.0, 5446.0, 6224.0, 7002.0]<br>2800, [-9801.0, -7623.0, -6534.0, -5445.0, -4356.0, -3267.0, -2178.0, -1089.0, -10.0, 10.0, 1089.0, 2178.0, 3267.0, 4356.0, 5445.0, 6534.0, 7623.0, 9801.0]<br>3600, [-12600.0, -11200.0, -9800.0, -7000.0, -5600.0, -4200.0, -2800.0, -1400.0, -10.0, 10.0, 1400.0, 2800.0, 4200.0, 5600.0, 7000.0, 9800.0, 11200.0, 12600.0] |
| | 15 °C<br>900 MHz | 150, [-528.0, -480.0, -432.0, -384.0, -336.0, -288.0, -240.0, -192.0, -144.0, -96.0, -48.0, -10.0, 10.0, 48.0, 96.0, 144.0, 192.0, 240.0, 288.0, 336.0, 384.0, 432.0, 480.0, 528.0]<br>800, [-2805.0, -2550.0, -2295.0, -2040.0, -1785.0, -1530.0, -1275.0, -1020.0, -765.0, -510.0, -255.0, -10.0, 10.0, 255.0, 510.0, 765.0, 1020.0, 1275.0, 1530.0, 1785.0, 2040.0, 2295.0, 2550.0, 2805.0]<br>1200, [-4202.0, -3820.0, -3438.0, -3056.0, -2674.0, -2292.0, -1910.0, -1528.0, -1146.0, -764.0, -382.0, -10.0, 10.0, 382.0, 764.0, 1146.0, 1528.0, 1910.0, 2292.0, 2674.0, 3056.0, 3438.0, 3820.0, 4202.0]<br>1600, [-5599.0, -5090.0, -4581.0, -4072.0, -3563.0, -3054.0, -2545.0, -2036.0, -1527.0, -1018.0, -509.0, -10.0, 10.0, 509.0, 1018.0, 1527.0, 2036.0, 2545.0, 3054.0, 3563.0, 4072.0, 4581.0, 5090.0, 5599.0]<br>2000, [-6996.0, -6360.0, -5724.0, -5088.0, -4452.0, -3816.0, -3180.0, -2544.0, -1908.0, -1272.0, -636.0, -10.0, 10.0, 636.0, 1272.0, 1908.0, 2544.0, 3180.0, 3816.0, 4452.0, 5088.0, 5724.0, 6360.0, 6996.0]<br>2400, [-7640.0, -6876.0, -6112.0, -5348.0, -4584.0, -3820.0, -3056.0, -2292.0, -1528.0, -764.0, -10.0, 10.0, 764.0, 1528.0, 2292.0, 3056.0, 3820.0, 4584.0, 5348.0, 6112.0, 6876.0, 7640.0] |

| | | |
|---|---|---|
| | | 2800, [-9801.0, -8019.0, -7128.0, -6237.0, -5346.0, -4455.0, -3564.0, -2673.0, -1782.0, -891.0, -10.0, 10.0, 891.0, 1782.0, 2673.0, 3564.0, 4455.0, 5346.0, 6237.0, 7128.0, 8019.0, 9801.0]<br>3600, [-12595.0, -11450.0, -10305.0, -6870.0, -5725.0, -4580.0, -3435.0, -2290.0, -1145.0, -10.0, 10.0, 1145.0, 2290.0, 3435.0, 4580.0, 5725.0, 6870.0, 10305.0, 11450.0, 12595.0] |
| | 20 °C<br>900 MHz | 200, [-704.0, -640.0, -576.0, -512.0, -448.0, -384.0, -320.0, -256.0, -192.0, -128.0, -64.0, -10.0, 10.0, 64.0, 128.0, 192.0, 256.0, 320.0, 384.0, 448.0, 512.0, 576.0, 640.0, 704.0]<br>500, [-1749.0, -1590.0, -1431.0, -1272.0, -1113.0, -954.0, -795.0, -636.0, -477.0, -318.0, -159.0, -10.0, 10.0, 159.0, 318.0, 477.0, 636.0, 795.0, 954.0, 1113.0, 1272.0, 1431.0, 1590.0, 1749.0]<br>1000, [-3498.0, -3180.0, -2862.0, -2544.0, -2226.0, -1908.0, -1590.0, -1272.0, -954.0, -636.0, -318.0, -10.0, 10.0, 318.0, 636.0, 954.0, 1272.0, 1590.0, 1908.0, 2226.0, 2544.0, 2862.0, 3180.0, 3498.0]<br>1500, [-5247.0, -4770.0, -4293.0, -3816.0, -3339.0, -2862.0, -2385.0, -1908.0, -1431.0, -954.0, -477.0, -10.0, 10.0, 477.0, 954.0, 1431.0, 1908.0, 2385.0, 2862.0, 3339.0, 3816.0, 4293.0, 4770.0, 5247.0]<br>2000, [-6996.0, -6360.0, -5724.0, -5088.0, -4452.0, -3816.0, -3180.0, -2544.0, -1908.0, -1272.0, -636.0, -10.0, 10.0, 636.0, 1272.0, 1908.0, 2544.0, 3180.0, 3816.0, 4452.0, 5088.0, 5724.0, 6360.0, 6996.0]<br>2500, [-7950.0, -7155.0, -6360.0, -5565.0, -4770.0, -3975.0, -3180.0, -2385.0, -1590.0, -795.0, -10.0, 10.0, 795.0, 1590.0, 2385.0, 3180.0, 3975.0, 4770.0, 5565.0, 6360.0, 7155.0, 7950.0] |

| | | |
|---|---|---|
| | | 3000, [-10505.0, -9550.0, -7640.0, -6685.0, -5730.0, -4775.0, -3820.0, -2865.0, -1910.0, -955.0, -10.0, 10.0, 955.0, 1910.0, 2865.0, 3820.0, 4775.0, 5730.0, 6685.0, 7640.0, 9550.0, 10505.0]<br>3600, [-12595.0, -11450.0, -10305.0, -9160.0, -6870.0, -5725.0, -4580.0, -3435.0, -2290.0, -1145.0, -10.0, 10.0, 1145.0, 2290.0, 3435.0, 4580.0, 5725.0, 6870.0, 9160.0, 10305.0, 11450.0, 12595.0] |
| **LE A12(C2)** | 20 °C<br>900 MHz | 200, [-704.0, -640.0, -576.0, -512.0, -448.0, -384.0, -320.0, -256.0, -192.0, -128.0, -64.0, -10.0, 10.0, 64.0, 128.0, 192.0, 256.0, 320.0, 384.0, 448.0, 512.0, 576.0, 640.0, 704.0]<br>500, [-1749.0, -1590.0, -1431.0, -1272.0, -1113.0, -954.0, -795.0, -636.0, -477.0, -318.0, -159.0, -10.0, 10.0, 159.0, 318.0, 477.0, 636.0, 795.0, 954.0, 1113.0, 1272.0, 1431.0, 1590.0, 1749.0]<br>1000, [-3498.0, -3180.0, -2862.0, -2544.0, -2226.0, -1908.0, -1590.0, -1272.0, -954.0, -636.0, -318.0, -10.0, 10.0, 318.0, 636.0, 954.0, 1272.0, 1590.0, 1908.0, 2226.0, 2544.0, 2862.0, 3180.0, 3498.0]<br>1500, [-5247.0, -4770.0, -4293.0, -3816.0, -3339.0, -2862.0, -2385.0, -1908.0, -1431.0, -954.0, -477.0, -10.0, 10.0, 477.0, 954.0, 1431.0, 1908.0, 2385.0, 2862.0, 3339.0, 3816.0, 4293.0, 4770.0, 5247.0]<br>2000, [-6996.0, -6360.0, -5724.0, -5088.0, -4452.0, -3816.0, -3180.0, -2544.0, -1908.0, -1272.0, -636.0, -10.0, 10.0, 636.0, 1272.0, 1908.0, 2544.0, 3180.0, 3816.0, 4452.0, 5088.0, 5724.0, 6360.0, 6996.0]<br>2500, [-7950.0, -7155.0, -6360.0, -5565.0, -4770.0, -3975.0, -3180.0, -2385.0, -1590.0, -795.0, -10.0, 10.0, 795.0, 1590.0, 2385.0, 3180.0, 3975.0, 4770.0, 5565.0, 6360.0, 7155.0, 7950.0] |

| | | |
|---|---|---|
| | | 3000, [-10505.0, -9550.0, -8595.0, -7640.0, -6685.0, -5730.0, -4775.0, -3820.0, -2865.0, -1910.0, -955.0, -10.0, 10.0, 955.0, 1910.0, 2865.0, 3820.0, 4775.0, 5730.0, 6685.0, 7640.0, 9550.0, 10505.0] |
| **LE T9(C6)** | 20 °C 900 MHz | 200, [-704.0, -640.0, -576.0, -512.0, -448.0, -384.0, -320.0, -256.0, -192.0, -128.0, -64.0, -10.0, 10.0, 64.0, 128.0, 192.0, 256.0, 320.0, 384.0, 448.0, 512.0, 576.0, 640.0, 704.0]<br>1000, [-3498.0, -3180.0, -2862.0, -2544.0, -2226.0, -1908.0, -1590.0, -1272.0, -954.0, -636.0, -318.0, -10.0, 10.0, 318.0, 636.0, 954.0, 1272.0, 1590.0, 1908.0, 2226.0, 2544.0, 2862.0, 3180.0, 3498.0]<br>1500, [-5247.0, -4770.0, -4293.0, -3816.0, -3339.0, -2862.0, -2385.0, -1908.0, -1431.0, -954.0, -477.0, -10.0, 10.0, 477.0, 954.0, 1431.0, 1908.0, 2385.0, 2862.0, 3339.0, 3816.0, 4293.0, 4770.0, 5247.0]<br>2000, [-6996.0, -6360.0, -5724.0, -5088.0, -4452.0, -3816.0, -3180.0, -2544.0, -1908.0, -1272.0, -636.0, -10.0, 10.0, 636.0, 1272.0, 1908.0, 2544.0, 3180.0, 3816.0, 4452.0, 5088.0, 5724.0, 6360.0, 6996.0]<br>2800, [-9801.0, -8019.0, -7128.0, -6237.0, -5346.0, -4455.0, -3564.0, -2673.0, -1782.0, -891.0, -10.0, 10.0, 891.0, 1782.0, 2673.0, 3564.0, 4455.0, 5346.0, 6237.0, 7128.0, 8019.0, 9801.0]<br>3600, [-12595.0, -11450.0, -10305.0, -8015.0, -6870.0, -5725.0, -4580.0, -3435.0, -2290.0, -1145.0, -10.0, 10.0, 1145.0, 2290.0, 3435.0, 4580.0, 5725.0, 6870.0, 8015.0, 9160.0, 10305.0, 11450.0, 12595.0] |
| **LE T9(C1')** | 20 °C 900 MHz | 200, [-704.0, -640.0, -576.0, -512.0, -448.0, -384.0, -320.0, -256.0, -192.0, -128.0, -64.0, -10.0, 10.0, 64.0, 128.0, 192.0, 256.0, 320.0, 384.0, 448.0, 512.0, 576.0, 640.0, 704.0] |

| | | |
|---|---|---|
| | | 2800, [-8910.0, -8019.0, -7128.0, -6237.0, -5346.0, -4455.0, -3564.0, -2673.0, -1782.0, -891.0, -10.0, 10.0, 891.0, 1782.0, 2673.0, 3564.0, 4455.0, 5346.0, 6237.0, 7128.0, 8019.0, 8910.0, 9801.0]<br>3600, [-12595.0, -11450.0, -10305.0, -9160.0, -8015.0, -6870.0, -5725.0, -4580.0, -3435.0, -2290.0, -1145.0, -10.0, 10.0, 1145.0, 2290.0, 3435.0, 4580.0, 5725.0, 6870.0, 8015.0, 9160.0, 10305.0, 11450.0, 12595.0] |

**Table S6.** Exchange parameters for LE, obtained from global or individual fitting the off-resonance $^{13}C$ $R_{1\rho}$ data to the Bloch-McConnell equation. Error bars represent experimental uncertainty (one s.d.) calculated by the Monte-Carlo approach from a single RD measurement (see Methods).

| **Residue** | **Parameter** | **7.5 °C, 900 MHz** | **10 °C, 900 MHz** | **15 °C, 900 MHz** | **20 °C, 900 MHz** |
|---|---|---|---|---|---|
| | | **Global Fit** | | | |
| **A12 (C8)** | $p_{minor,C8}$ (%) | 15 ± 1 | 15 ± 2 | 13 ± 2 | 9 ± 2 |
| | $k_{ex,C8}$ ($s^{-1}$) | 2100 ± 700 | 27000 ± 1000 | 42000 ± 2000 | 57000 ± 5000 |
| | $R_{1,C8}$ (Hz) | 2.9 ± 0.7 | 1.9 ± 0.5 | 1.9 ± 0.2 | 1.9 ± 0.1 |
| | $R_{2,C8}$ (Hz) | 40 ± 4 | 37 ± 7 | 41 ± 6 | 43 ± 5 |
| | $\Delta\omega_{C8}$ (ppm) | 3.8 ± 0.1 | | | |

| **Residue** | **Parameter** | **7.5 °C, 900 MHz** | **10 °C, 900 MHz** | **15 °C, 900 MHz** | **20 °C, 900 MHz** |
|---|---|---|---|---|---|
| | | **Individual Fit** | | | |
| **A12 (C8)** | $p_{minor,C8}$ (%) | 14 ± 3 | 16 ± 2 | 14 ± 2 | 8 ± 2 |
| | $k_{ex,C8}$ ($s^{-1}$) | 21000 ± 1000 | 27000 ± 800 | 41000 ± 2000 | 57000 ± 5000 |
| | $R_{1,C8}$ (Hz) | 2.9 ± 0.5 | 1.9 ± 0.3 | 1.9 ± 0.1 | 1.9 ± 0.1 |
| | $R_{2,C8}$ (Hz) | 40 ± 7 | 37 ± 4 | 43 ± 4 | 42 ± 5 |
| | $\Delta\omega_{C8}$ (ppm) | 3.9 ± 0.2 | 3.8 ± 0.2 | 3.6 ± 0.2 | 4.1 ± 0.4 |

**Table S7.** Full list of 63 RNA A-U bps adjacent to the dinucleotide apical loop present in PDB crystal structures.

| PDB ID | Nucleotide 1 | Nucleotide 2 | Base Pair | LW | Conformation 1 | Conformation 2 | C1'-C1' distance. Å | Hydrogen bonds | Bp conformation |
|---|---|---|---|---|---|---|---|---|---|
| 1vqm | 1:0U734 | 1:0.A737 | U-A | tSH | anti | anti | 8.241 | O2(carbonyl)-N6(amino)[3.74] | Other |
| 1vqn | 1:0.U734 | 1:0.A737 | U-A | tSH | anti | anti | 8.278 | O2(carbonyl)-N6(amino)[3.71] | Other |
| 1vql | 1:0.U734 | 1:0.A737 | U-A | tSH | anti | anti | 8.309 | O2(carbonyl)-N6(amino)[3.84] | Other |
| 1vqp | 1:0.U734 | 1:0.A737 | U-A | tSH | anti | anti | 8.401 | O2(carbonyl)-N6(amino)[3.91] | Other |
| 6chr | 1:A.A221 | 1:A.U224 | A-U | cWW | anti | anti | 8.953 | N6(amino)-O4(carbonyl)[3.84]<br>N1-N3(imino)[2.66] | Other |
| 6n5s | 1:A.A15 | 1:A.U18 | A-U | tSH | anti | anti | 9.03 | O2'(hydroxyl)-O4(carbonyl)[3.04] | Other |
| 4v89 | 1:BA.U2796 | 1:BA.A2799 | U-A | cWW | anti | anti | 9.512 | N3(imino)-N1[2.87] | Other |
| 4v85 | 1:BA.U2796 | 1:BA.A2799 | U-A | cWW | anti | anti | 9.574 | N3(imino)-N1[3.12] | Other |
| 4v9c | 1:BA.U2796 | 1:BA.A2799 | U-A | cWW | anti | -- | 9.942 | N3(imino)-N1[2.43]<br>O4(carbonyl)-N6(amino)[2.86] | Other |
| 5u31 | 1:B.U4 | 1:B.A7 | U-A | cWW | anti | anti | 10.001 | N3(imino)-N1[2.74]<br>O4(carbonyl)-N6(amino)[3.20] | Watson-Crick |
| 5u33 | 1:B.U4 | 1:B.A7 | U-A | cWW | anti | anti | 10.001 | N3(imino)-N1[2.85]<br>O4(carbonyl)-N6(amino)[3.48] | Watson-Crick |
| 4v9p | 1:GA.U2796 | 1:GA.A2799 | U-A | cWW | anti | -- | 10.136 | N3(imino)-N1[2.75]<br>O4(carbonyl)-N6(amino)[2.93] | Other |
| 4v9p | 1:AA.U2796 | 1:AA.A2799 | U-A | cWW | anti | -- | 10.183 | N3(imino)-N1[2.82]<br>O4(carbonyl)-N6(amino)[2.95] | Other |
| 4v9p | 1:EA.U2796 | 1:EA.A2799 | U-A | cWW | anti | -- | 10.303 | N3(imino)-N1[3.12]<br>O4(carbonyl)-N6(amino)[3.36] | Other |
| 5u30 | 1:B.U4 | 1:B.A7 | U-A | cWW | anti | anti | 10.31 | N3(imino)-N1[3.09]<br>O4(carbonyl)-N6(amino)[3.62] | Watson-Crick |
| 6m0x | 1:B.U56 | 1:B.A59 | U-A | cWW | anti | anti | 10.344 | N3(imino)-N1[2.93]<br>O4(carbonyl)-N6(amino)[3.41] | Watson-Crick |
| 2f8k | 1:B.U6 | 1:B.A9 | U-A | cWW | anti | anti | 10.358 | N3(imino)-N1[2.70]<br>O4(carbonyl)-N6(amino)[2.93] | Watson-Crick |
| 6m0w | 1:B.U56 | 1:B.A59 | U-A | cWW | anti | anti | 10.358 | N3(imino)-N1[2.87]<br>O4(carbonyl)-N6(amino)[3.32] | Watson-Crick |
| 5u34 | 1:B.U4 | 1:B.A7 | U-A | cWW | anti | anti | 10.362 | N3(imino)-N1[3.11]<br>O4(carbonyl)-N6(amino)[3.62] | Watson-Crick |
| 4v9o | 1:CA.U2796 | 1:CA.A2799 | U-A | cWW | anti | -- | 10.374 | N3(imino)-N1[2.96]<br>O4(carbonyl)-N6(amino)[3.11] | Other |
| 6m0v | 1:B.U56 | 1:B.A59 | U-A | cWW | anti | anti | 10.388 | N3(imino)-N1[3.21]<br>O4(carbonyl)-N6(amino)[3.86] | Watson-Crick |

| | | | | | | | | | |
|---|---|---|---|---|---|---|---|---|---|
| 4u27 | 1:DA.U2796 | 1:DA.A2799 | U-A | cWW | anti | -- | 10.411 | N3(imino)-N1[3.30] | Other |
| 4v8p | 1:A1.U2558 | 1:A1.A2561 | U-A | cWW | anti | anti | 10.483 | N3(imino)-N1[3.43]<br>O4(carbonyl)-N6(amino)[3.97] | Watson-Crick |
| 4v8p | 1:F1.U2558 | 1:F1.A2561 | U-A | cWW | anti | anti | 10.488 | N3(imino)-N1[3.42]<br>O4(carbonyl)-N6(amino)[3.98] | Watson-Crick |
| 4v8p | 1:H1.U2558 | 1:H1.A2561 | U-A | cWW | anti | anti | 10.493 | N3(imino)-N1[3.42]<br>O4(carbonyl)-N6(amino)[3.96] | Watson-Crick |
| 4v8p | 1:D1.U2558 | 1:D1.A2561 | U-A | cWW | anti | anti | 10.499 | N3(imino)-N1[3.45]<br>O4(carbonyl)-N6(amino)[3.99] | Watson-Crick |
| 4v9o | 1:EA.U2796 | 1:EA.A2799 | U-A | cWW | anti | -- | 10.511 | N3(imino)-N1[2.86]<br>O4(carbonyl)-N6(amino)[2.89] | Other |
| 4v7u | 1:BA.U2796 | 1:BA.A2799 | U-A | cWW | anti | -- | 10.554 | N3(imino)-N1[3.07]<br>O4(carbonyl)-N6(amino)[3.10] | Other |
| 5wqe | 1:B.U4 | 1:B.A7 | U-A | cWW | anti | anti | 10.557 | N3(imino)-N1[2.66]<br>O4(carbonyl)-N6(amino)[2.49] | Watson-Crick |
| 6i7v | 1:DA.U2796 | 1:DA.A2799 | U-A | cWW | anti | -- | 10.628 | N3(imino)-N1[3.08] | Other |
| 4v7v | 1:BA.U2796 | 1:BA.A2799 | U-A | cWW | anti | -- | 10.644 | N3(imino)-N1[3.03]<br>O4(carbonyl)-N6(amino)[3.09] | Other |
| 4v9o | 1:GA.U2796 | 1:GA.A2799 | U-A | cWW | anti | -- | 10.679 | N3(imino)-N1[3.08]<br>O4(carbonyl)-N6(amino)[3.26] | Other |
| 4v9p | 1:CA.U2796 | 1:CA.A2799 | U-A | cWW | anti | -- | 10.697 | N3(imino)-N1[2.87]<br>O4(carbonyl)-N6(amino)[2.65] | Other |
| 4u1v | 1:BA.U2796 | 1:BA.A2799 | U-A | cWW | anti | -- | 10.754 | N3(imino)-N1[3.13]<br>O4(carbonyl)-N6(amino)[3.06] | Other |
| 4v9d | 1:DA.U2796 | 1:DA.A2799 | U-A | cWW | anti | -- | 10.758 | O2(carbonyl)*N1[3.20]<br>N3(imino)-N1[3.37] | Other |
| 4u1u | 1:BA.U2796 | 1:BA.A2799 | U-A | cWW | anti | -- | 10.784 | N3(imino)-N1[2.90]<br>O4(carbonyl)-N6(amino)[2.63] | Other |
| 4u26 | 1:BA.U2796 | 1:BA.A2799 | U-A | cWW | anti | -- | 10.79 | N3(imino)-N1[3.07]<br>O4(carbonyl)-N6(amino)[2.96] | Other |
| 4v7s | 1:BA.U2796 | 1:BA.A2799 | U-A | cWW | anti | -- | 10.796 | N3(imino)-N1[3.05]<br>O4(carbonyl)-N6(amino)[2.85] | Other |
| 4u24 | 1:BA.U2796 | 1:BA.A2799 | U-A | cWW | anti | -- | 10.823 | N3(imino)-N1[3.03]<br>O4(carbonyl)-N6(amino)[2.78] | Other |
| 6by1 | 1:CA.U2796 | 1:CA.A2799 | U-A | cWW | anti | -- | 10.846 | N3(imino)-N1[2.84]<br>O4(carbonyl)-N6(amino)[2.73] | Other |
| 6by1 | 1:DA.U2796 | 1:DA.A2799 | U-A | cWW | anti | -- | 10.872 | N3(imino)-N1[2.76]<br>O4(carbonyl)-N6(amino)[2.71] | Other |
| 4v7t | 1:BA.U2796 | 1:BA.A2799 | U-A | cWW | anti | -- | 10.886 | N3(imino)-N1[3.23]<br>O4(carbonyl)-N6(amino)[2.94] | Other |
| 5it8 | 1:DA.U2796 | 1:DA.A2799 | U-A | cWW | anti | -- | 10.886 | O2(carbonyl)*N1[3.11]<br>N3(imino)-N1[3.34] | Other |

| | | | | | | | | | |
|---|---|---|---|---|---|---|---|---|---|
| | | | | | | | | O4(carbonyl)-N6(amino)[3.13] | |
| 4v6c | 1:BA.U2796 | 1:BA.A2799 | U-A | cWW | anti | -- | 10.889 | N3(imino)-N1[3.23]<br>O4(carbonyl)-N6(amino)[3.41] | Other |
| 4v9d | 1:CA.U2796 | 1:CA.A2799 | U-A | cWW | anti | -- | 10.903 | N3(imino)-N1[3.27]<br>O4(carbonyl)-N6(amino)[3.17] | Other |
| 5it8 | 1:CA.U2796 | 1:CA.A2799 | U-A | cWW | anti | -- | 10.903 | O2(carbonyl)*N1[3.11]<br>N3(imino)-N1[3.32]<br>O4(carbonyl)-N6(amino)[3.12] | Other |
| 4u20 | 1:BA.U2796 | 1:BA.A2799 | U-A | cWW | anti | -- | 10.922 | N3(imino)-N1[3.05]<br>O4(carbonyl)-N6(amino)[2.90] | Other |
| 5j5b | 1:CA.U2796 | 1:CA.A2799 | U-A | cWW | anti | -- | 10.938 | N3(imino)-N1[3.20]<br>O4(carbonyl)-N6(amino)[2.79] | Other |
| 5j91 | 1:CA.U2796 | 1:CA.A2799 | U-A | cWW | anti | -- | 10.938 | N3(imino)-N1[3.20]<br>O4(carbonyl)-N6(amino)[2.79] | Other |
| 5j5b | 1:DA.U2796 | 1:DA.A2799 | U-A | cWW | anti | -- | 10.953 | N3(imino)-N1[3.21]<br>O4(carbonyl)-N6(amino)[2.78] | Other |
| 5j91 | 1:DA.U2796 | 1:DA.A2799 | U-A | cWW | anti | -- | 10.953 | N3(imino)-N1[3.21]<br>O4(carbonyl)-N6(amino)[2.78] | Other |
| 4u25 | 1:BA.U2796 | 1:BA.A2799 | U-A | cWW | anti | -- | 10.996 | N3(imino)-N1[3.14]<br>O4(carbonyl)-N6(amino)[2.95] | Other |
| 5j8a | 1:CA.U2796 | 1:CA.A2799 | U-A | cWW | anti | -- | 10.998 | N3(imino)-N1[3.30]<br>O4(carbonyl)-N6(amino)[2.92] | Other |
| 5j7l | 1:CA.U2796 | 1:CA.A2799 | U-A | cWW | anti | -- | 11.013 | O2(carbonyl)*N1[3.09]<br>N3(imino)-N1[3.31]<br>O4(carbonyl)-N6(amino)[2.87] | Other |
| 5j8a | 1:DA.U2796 | 1:DA.A2799 | U-A | cWW | anti | -- | 11.033 | O2(carbonyl)*N1[3.10]<br>N3(imino)-N1[3.32]<br>O4(carbonyl)-N6(amino)[2.92] | Other |
| 5j7l | 1:DA.U2796 | 1:DA.A2799 | U-A | cWW | anti | -- | 11.041 | O2(carbonyl)*N1[3.09]<br>N3(imino)-N1[3.32]<br>O4(carbonyl)-N6(amino)[2.86] | Other |
| 4wf1 | 1:BA.U2796 | 1:BA.A2799 | U-A | cWW | anti | -- | 11.12 | N3(imino)-N1[3.12]<br>O4(carbonyl)-N6(amino)[2.82] | Other |
| 4v9o | 1:AA.U2796 | 1:AA.A2799 | U-A | cWW | anti | -- | 11.241 | N3(imino)-N1[3.04]<br>O4(carbonyl)-N6(amino)[2.43] | Other |
| 4u27 | 1:BA.U2796 | 1:BA.A2799 | U-A | cWW | anti | -- | 11.248 | N3(imino)-N1[3.26]<br>O4(carbonyl)-N6(amino)[2.95] | Other |
| 4ybb | 1:DA.U2796 | 1:DA.A2799 | U-A | cWW | anti | -- | 11.426 | N3(imino)-N1[3.24]<br>O4(carbonyl)-N6(amino)[2.50] | Other |
| 4v4z | 1:BA.U2702 | 1:BA.A2705 | U-A | c.H | anti | anti | 12.444 | O4(carbonyl)-N6(amino)[2.26] | Other |
| 4v4y | 1:BA.U2702 | 1:BA.A2705 | U-A | cHH | anti | anti | 12.597 | O4(carbonyl)-N6(amino)[2.87] | Other |

| | | | | | | | | | |
|---|---|---|---|---|---|---|---|---|---|
| 4v4x | 1:BA.U2702 | 1:BA.A2705 | U-A | c.W | anti | anti | 12.607 | O4(carbonyl)-N6(amino)[2.34] | Other |

**Table S8.** Full list of 228 RNA A-U bps adjacent to the trinucleotide apical loop present in PDB crystal structures.

| PDB ID | Nucleotide 1 | Nucleotide 2 | Base Pair | LW | Conformation 1 | Conformation 2 | C1'-C1' distance, Å | Hydrogen bonds | Bp conformation |
|---|---|---|---|---|---|---|---|---|---|
| 1fjg | 1:A.U81 | 1:A.A88 | U-A | t.H | anti | anti | 9.458 | O2(carbonyl)-N6(amino)[2.88] | Other |
| 1gid | 1:B.A235 | 1:B.U239 | A-U | cWW | anti | anti | 10.829 | N6(amino)-O4(carbonyl)[2.32]<br>N1-N3(imino)[2.78] | Watson-Crick |
| 1gid | 1:A.A235 | 1:A.U239 | A-U | cWW | anti | anti | 10.83 | N6(amino)-O4(carbonyl)[2.31]<br>N1-N3(imino)[2.77] | Watson-Crick |
| 1hnw | 1:A.U81 | 1:A.A88 | U-A | tSH | anti | anti | 9.792 | O2(carbonyl)-N6(amino)[3.72] | Other |
| 1hnx | 1:A.U81 | 1:A.A88 | U-A | tSH | anti | anti | 9.979 | O2(carbonyl)-N6(amino)[3.66] | Other |
| 1hnz | 1:A.U81 | 1:A.A88 | U-A | t.H | anti | anti | 9.624 | O2(carbonyl)-N6(amino)[3.00] | Other |
| 1hr0 | 1:A.U81 | 1:A.A88 | U-A | tSH | anti | anti | 9.563 | O2(carbonyl)-N6(amino)[3.21] | Other |
| 1hr2 | 1:B.A235 | 1:B.U239 | A-U | cWW | anti | anti | 11.335 | N6(amino)-O4(carbonyl)[2.92]<br>N1-N3(imino)[3.23] | Other |
| 1hr2 | 1:A.A235 | 1:A.U239 | A-U | cWW | anti | anti | 11.398 | N6(amino)-O4(carbonyl)[3.04]<br>N1-N3(imino)[3.44] | Other |
| 1i94 | 1:A.U79 | 1:A.A83 | U-A | tWH | anti | anti | 11.44 | O2(carbonyl)-N6(amino)[2.78] | Other |
| 1i95 | 1:A.U79 | 1:A.A83 | U-A | tWH | anti | anti | 9.49 | O2(carbonyl)-N6(amino)[2.71]<br>N3(imino)-N7[2.68] | Other |
| 1i96 | 1:A.U79 | 1:A.A83 | U-A | tWH | anti | anti | 11.569 | O2(carbonyl)-N6(amino)[2.91] | Other |
| 1ibk | 1:A.U81 | 1:A.A88 | U-A | tSH | anti | anti | 9.528 | O2(carbonyl)-N6(amino)[2.98] | Other |
| 1j5e | 1:A.U81 | 1:A.A88 | U-A | t.H | anti | anti | 9.746 | O2(carbonyl)-N6(amino)[3.19] | Other |
| 1l8v | 1:A.A235 | 1:A.U239 | A-U | cWW | anti | anti | 10.188 | N1-N3(imino)[3.17] | Other |
| 1l8v | 1:B.A235 | 1:B.U239 | A-U | cWW | anti | anti | 10.19 | N1-N3(imino)[3.18] | Other |
| 1n33 | 1:A.U81 | 1:A.A88 | U-A | tWH | anti | anti | 10.569 | O2(carbonyl)-N6(amino)[3.22] | Other |
| 2e5l | 1:A.U81 | 1:A.A88 | U-A | tWH | anti | anti | 10.112 | O2(carbonyl)-N6(amino)[2.56] | Other |
| 2hhh | 1:A.U80 | 1:A.A84 | U-A | tWH | anti | anti | 10.072 | O2(carbonyl)-N6(amino)[3.17] | Other |
| 2zm6 | 1:A.U81 | 1:A.A88 | U-A | tWH | anti | anti | 10.334 | O2(carbonyl)-N6(amino)[2.81] | Other |
| 4dr2 | 1:A.U81 | 1:A.A88 | U-A | tSH | anti | anti | 9.998 | O2(carbonyl)-N6(amino)[3.85] | Other |
| 4dr5 | 1:A.U81 | 1:A.A88 | U-A | tWH | anti | anti | 10.847 | O2(carbonyl)-N6(amino)[3.28] | Other |
| 4duy | 1:A.U81 | 1:A.A88 | U-A | tSH | anti | anti | 9.812 | O2(carbonyl)-N6(amino)[3.71] | Other |
| 4lf4 | 1:A.U81 | 1:A.A88 | U-A | t.H | anti | anti | 9.526 | O2(carbonyl)-N6(amino)[2.39] | Other |

| | | | | | | | | | |
|---|---|---|---|---|---|---|---|---|---|
| 4lf5 | 1:A.U81 | 1:A.A88 | U-A | tWH | anti | anti | 9.707 | O2(carbonyl)-N6(amino)[2.59] | Other |
| 4lf6 | 1:A.U81 | 1:A.A88 | U-A | tWH | anti | anti | 10.13 | O2(carbonyl)-N6(amino)[2.28] | Other |
| 4lf7 | 1:A.U81 | 1:A.A88 | U-A | t.H | anti | anti | 9.922 | O2(carbonyl)-N6(amino)[2.44] | Other |
| 4lf8 | 1:A.U81 | 1:A.A88 | U-A | t.H | anti | anti | 9.922 | O2(carbonyl)-N6(amino)[2.44] | Other |
| 4lf9 | 1:A.U81 | 1:A.A88 | U-A | tSH | anti | anti | 9.618 | O2(carbonyl)-N6(amino)[3.28] | Other |
| 4lfa | 1:A.U81 | 1:A.A88 | U-A | tSH | anti | anti | 9.537 | O2(carbonyl)-N6(amino)[3.60] | Other |
| 4lfc | 1:A.U81 | 1:A.A88 | U-A | t.H | anti | anti | 9.593 | O2(carbonyl)-N6(amino)[2.86] | Other |
| 4ox9 | 1:A.U81 | 1:A.A88 | U-A | t.H | anti | anti | 9.746 | O2(carbonyl)-N6(amino)[3.19] | Other |
| 4p8z | 1:A.U128 | 1:A.A132 | U-A | tWH | anti | anti | 11.017 | O2(carbonyl)-N6(amino)[3.46] | Other |
| 4u3m | 1:5.U1567 | 1:5.A1571 | U-A | cWW | anti | anti | 9.746 | N3(imino)-N1[2.95] | Other |
| 4u3n | 1:6.U1397 | 1:6.A1401 | U-A | tWH | anti | anti | 9.703 | O2(carbonyl)-N6(amino)[3.66] | Other |
| 4u3n | 1:5.U1567 | 1:5.A1571 | U-A | cWW | anti | anti | 9.869 | N3(imino)-N1[3.45] | Other |
| 4u3n | 1:2.U1397 | 1:2.A1401 | U-A | tWH | anti | anti | 10.155 | O2(carbonyl)-N6(amino)[3.38] | Other |
| 4u3u | 1:6.U1397 | 1:6.A1401 | U-A | tWH | anti | anti | 9.934 | O2(carbonyl)-N6(amino)[3.13] | Other |
| 4u3u | 1:5.U1567 | 1:5.A1571 | U-A | cWW | anti | anti | 9.983 | N3(imino)-N1[3.32] | Other |
| 4u3u | 1:2.U1397 | 1:2.A1401 | U-A | tWH | anti | anti | 10.252 | O2(carbonyl)-N6(amino)[3.41] | Other |
| 4u4n | 1:5.U1567 | 1:5.A1571 | U-A | cWW | anti | anti | 9.761 | N3(imino)-N1[3.09] | Other |
| 4u4n | 1:2.U1397 | 1:2.A1401 | U-A | t.H | anti | anti | 10.147 | O2'(hydroxyl)-N6(amino)[3.65] O2(carbonyl)-N6(amino)[3.76] | Other |
| 4u4o | 1:5.U1567 | 1:5.A1571 | U-A | cWW | anti | -- | 9.47 | N3(imino)-N1[3.04] | Other |
| 4u4o | 1:2.U1397 | 1:2.A1401 | U-A | tWH | anti | anti | 10.126 | O2(carbonyl)-N6(amino)[2.50] | Other |
| 4u4o | 1:6.U1397 | 1:6.A1401 | U-A | tWH | anti | anti | 10.176 | O2(carbonyl)-N6(amino)[3.67] | Other |
| 4u4q | 1:6.U1397 | 1:6.A1401 | U-A | tWH | anti | anti | 9.455 | O2(carbonyl)-N6(amino)[3.87] | Other |
| 4u4q | 1:5.U1567 | 1:5.A1571 | U-A | cWW | anti | anti | 9.826 | N3(imino)-N1[3.37] | Other |
| 4u4q | 1:2.U1397 | 1:2.A1401 | U-A | t.H | anti | anti | 10.385 | O2(carbonyl)-N6(amino)[3.67] | Other |
| 4u4r | 1:5.U1567 | 1:5.A1571 | U-A | cWW | anti | anti | 9.722 | N3(imino)-N1[2.88] O4(carbonyl)-N6(amino)[3.75] | Other |
| 4u4r | 1:6.U1397 | 1:6.A1401 | U-A | tWH | anti | anti | 9.762 | O2(carbonyl)-N6(amino)[3.53] | Other |
| 4u4r | 1:2.U1397 | 1:2.A1401 | U-A | tWH | anti | anti | 10.415 | O2(carbonyl)-N6(amino)[3.85] | Other |
| 4u4u | 1:5.U1567 | 1:5.A1571 | U-A | cWW | anti | anti | 9.737 | N3(imino)-N1[3.28] | Other |

| | | | | | | | | | |
|---|---|---|---|---|---|---|---|---|---|
| 4u4u | 1:2.U1397 | 1:2.A1401 | U-A | t.H | anti | anti | 10.414 | O2(carbonyl)-N6(amino)[3.72] | Other |
| 4u4y | 1:5.U1567 | 1:5.A1571 | U-A | cWW | anti | anti | 9.834 | N3(imino)-N1[3.41] | Other |
| 4u4y | 1:2.U1397 | 1:2.A1401 | U-A | tWH | anti | anti | 10.407 | O2(carbonyl)-N6(amino)[3.40] | Other |
| 4u4z | 1:1.U1567 | 1:1.A1571 | U-A | cSW | anti | anti | 7.828 | O2(carbonyl)-N6(amino)[2.10] | Other |
| 4u4z | 1:2.U1397 | 1:2.A1401 | U-A | tWH | anti | anti | 9.985 | O2(carbonyl)-N6(amino)[3.30] | Other |
| 4u50 | 1:5.U1567 | 1:5.A1571 | U-A | cWW | anti | anti | 9.842 | N3(imino)-N1[3.40] | Other |
| 4u50 | 1:2.U1397 | 1:2.A1401 | U-A | t.H | anti | anti | 10.09 | O2'(hydroxyl)-N6(amino)[3.58]<br>O2(carbonyl)-N6(amino)[3.73] | Other |
| 4u51 | 1:6.U1397 | 1:6.A1401 | U-A | tWH | anti | anti | 9.486 | O2(carbonyl)-N6(amino)[3.71] | Other |
| 4u51 | 1:5.U1567 | 1:5.A1571 | U-A | cWW | anti | anti | 9.646 | N3(imino)-N1[3.09] | Other |
| 4u51 | 1:2.U1397 | 1:2.A1401 | U-A | t.H | anti | anti | 10.128 | O2'(hydroxyl)-N6(amino)[3.68]<br>O2(carbonyl)-N6(amino)[3.68] | Other |
| 4u52 | 1:5.U1567 | 1:5.A1571 | U-A | cWW | anti | anti | 9.922 | N3(imino)-N1[3.17]<br>O4(carbonyl)-N6(amino)[3.95] | Other |
| 4u53 | 1:6.U1397 | 1:6.A1401 | U-A | tWH | anti | anti | 9.648 | O2(carbonyl)-N6(amino)[3.97] | Other |
| 4u53 | 1:5.U1567 | 1:5.A1571 | U-A | cWW | anti | anti | 9.913 | N3(imino)-N1[3.61] | Other |
| 4u53 | 1:2.U1397 | 1:2.A1401 | U-A | t.H | anti | anti | 10.465 | O2(carbonyl)-N6(amino)[3.50] | Other |
| 4u55 | 1:6.U1397 | 1:6.A1401 | U-A | tWH | anti | anti | 9.614 | O2(carbonyl)-N6(amino)[3.59] | Other |
| 4u55 | 1:5.U1567 | 1:5.A1571 | U-A | cWW | anti | anti | 9.891 | N3(imino)-N1[3.21] | Other |
| 4u56 | 1:1.U1567 | 1:1.A1571 | U-A | cSH | anti | anti | 8.223 | O2(carbonyl)-N6(amino)[2.09] | Other |
| 4u56 | 1:6.U1397 | 1:6.A1401 | U-A | tWH | anti | anti | 9.721 | O2(carbonyl)-N6(amino)[3.27] | Other |
| 4u56 | 1:5.U1567 | 1:5.A1571 | U-A | cWW | anti | anti | 9.993 | N3(imino)-N1[3.05]<br>O4(carbonyl)-N6(amino)[3.92] | Other |
| 4u6f | 1:6.U1397 | 1:6.A1401 | U-A | tWH | anti | anti | 9.697 | O2(carbonyl)-N6(amino)[3.59] | Other |
| 4u6f | 1:5.U1567 | 1:5.A1571 | U-A | cWW | anti | anti | 9.884 | N3(imino)-N1[3.27] | Other |
| 4u6f | 1:2.U1397 | 1:2.A1401 | U-A | tWH | anti | anti | 10.148 | O2(carbonyl)-N6(amino)[3.69] | Other |
| 4v4h | 1:DB.U2796 | 1:DB.A2800 | U-A | cWW | anti | anti | 10.717 | N3(imino)-N1[2.85]<br>O4(carbonyl)-N6(amino)[2.84] | Watson-Crick |
| 4v4h | 1:BB.U2796 | 1:BB.A2800 | U-A | cWW | anti | anti | 10.721 | N3(imino)-N1[2.85]<br>O4(carbonyl)-N6(amino)[2.84] | Watson-Crick |
| 4v4i | 1:y.U82 | 1:y.A87 | U-A | tWH | anti | anti | 9.89 | O2(carbonyl)-N6(amino)[2.28] | Other |
| 4v4q | 1:BB.U2796 | 1:BB.A2800 | U-A | cWW | anti | anti | 10.548 | N3(imino)-N1[2.75]<br>O4(carbonyl)-N6(amino)[2.84] | Watson-Crick |

| | | | | | | | | | |
|---|---|---|---|---|---|---|---|---|---|
| 4v4q | 1:DB.U2796 | 1:DB.A2800 | U-A | cWW | anti | anti | 10.561 | N3(imino)-N1[2.76]<br>O4(carbonyl)-N6(amino)[2.84] | Watson-Crick |
| 4v4x | 1:AA.U82 | 1:AA.A87 | U-A | cWW | anti | anti | 7.284 | N3(imino)-N1[2.73]<br>O4(carbonyl)-N6(amino)[3.01] | Other |
| 4v4x | 1:BA.U1082 | 1:BA.A1086 | U-A | cWW | anti | anti | 8.18 | N3(imino)-N1[2.76]<br>O4(carbonyl)-N6(amino)[2.97] | Other |
| 4v4y | 1:AA.U82 | 1:AA.A87 | U-A | cWW | anti | anti | 6.948 | N3(imino)-N1[2.75]<br>O4(carbonyl)-N6(amino)[3.00] | Other |
| 4v4y | 1:BA.U1082 | 1:BA.A1086 | U-A | cWW | anti | anti | 8.567 | N3(imino)-N1[2.75]<br>O4(carbonyl)-N6(amino)[2.98] | Other |
| 4v4z | 1:AA.U82 | 1:AA.A87 | U-A | cWW | anti | anti | 7.551 | N3(imino)-N1[2.69]<br>O4(carbonyl)-N6(amino)[3.03] | Other |
| 4v4z | 1:BA.U1082 | 1:BA.A1086 | U-A | cWW | anti | anti | 8.526 | N3(imino)-N1[2.77]<br>O4(carbonyl)-N6(amino)[2.99] | Other |
| 4v50 | 1:BB.U2796 | 1:BB.A2800 | U-A | cWW | anti | anti | 10.808 | N3(imino)-N1[2.92]<br>O4(carbonyl)-N6(amino)[2.82] | Watson-Crick |
| 4v50 | 1:DB.U2796 | 1:DB.A2800 | U-A | cWW | anti | anti | 10.816 | N3(imino)-N1[2.90]<br>O4(carbonyl)-N6(amino)[2.79] | Watson-Crick |
| 4v52 | 1:BB.U2796 | 1:BB.A2800 | U-A | cWW | anti | anti | 11.043 | N3(imino)-N1[2.99]<br>O4(carbonyl)-N6(amino)[2.74] | Other |
| 4v52 | 1:DB.U2796 | 1:DB.A2800 | U-A | cWW | anti | anti | 11.056 | N3(imino)-N1[3.01]<br>O4(carbonyl)-N6(amino)[2.75] | Other |
| 4v53 | 1:BB.U2796 | 1:BB.A2800 | U-A | cWW | anti | anti | 10.741 | N3(imino)-N1[2.91]<br>O4(carbonyl)-N6(amino)[2.91] | Watson-Crick |
| 4v53 | 1:DB.U2796 | 1:DB.A2800 | U-A | cWW | anti | anti | 10.744 | N3(imino)-N1[2.92]<br>O4(carbonyl)-N6(amino)[2.91] | Watson-Crick |
| 4v54 | 1:DB.U2796 | 1:DB.A2800 | U-A | cWW | anti | anti | 10.788 | N3(imino)-N1[2.88]<br>O4(carbonyl)-N6(amino)[2.77] | Watson-Crick |
| 4v54 | 1:BB.U2796 | 1:BB.A2800 | U-A | cWW | anti | anti | 10.795 | N3(imino)-N1[2.89]<br>O4(carbonyl)-N6(amino)[2.78] | Watson-Crick |
| 4v55 | 1:DB.U2796 | 1:DB.A2800 | U-A | cWW | anti | anti | 11.089 | N3(imino)-N1[3.09]<br>O4(carbonyl)-N6(amino)[2.84] | Other |
| 4v55 | 1:BB.U2796 | 1:BB.A2800 | U-A | cWW | anti | anti | 11.172 | N3(imino)-N1[3.08]<br>O4(carbonyl)-N6(amino)[2.78] | Other |
| 4v56 | 1:BB.U2796 | 1:BB.A2800 | U-A | cWW | anti | anti | 11.058 | N3(imino)-N1[2.98]<br>O4(carbonyl)-N6(amino)[2.68] | Other |
| 4v56 | 1:DB.U2796 | 1:DB.A2800 | U-A | cWW | anti | anti | 11.063 | N3(imino)-N1[2.99]<br>O4(carbonyl)-N6(amino)[2.69] | Other |
| 4v57 | 1:BB.U2796 | 1:BB.A2800 | U-A | cWW | anti | anti | 10.717 | N3(imino)-N1[2.85]<br>O4(carbonyl)-N6(amino)[2.79] | Watson-Crick |
| 4v57 | 1:DB.U2796 | 1:DB.A2800 | U-A | cWW | anti | anti | 10.727 | N3(imino)-N1[2.87]<br>O4(carbonyl)-N6(amino)[2.83] | Watson-Crick |

| | | | | | | | | | |
|---|---|---|---|---|---|---|---|---|---|
| 4v5b | 1:CB.U138 | 1:CB.A142 | U-A | cWW | anti | anti | 9.13 | N3(imino)-N1[3.45] | Other |
| 4v5y | 1:DB.U2796 | 1:DB.A2800 | U-A | cWW | anti | anti | 11.061 | N3(imino)-N1[3.00]<br>O4(carbonyl)-N6(amino)[2.71] | Other |
| 4v5y | 1:BB.U2796 | 1:BB.A2800 | U-A | cWW | anti | anti | 11.063 | N3(imino)-N1[2.99]<br>O4(carbonyl)-N6(amino)[2.70] | Other |
| 4v64 | 1:BB.U2796 | 1:BB.A2800 | U-A | cWW | anti | anti | 10.939 | N3(imino)-N1[2.98]<br>O4(carbonyl)-N6(amino)[2.82] | Other |
| 4v64 | 1:DB.U2796 | 1:DB.A2800 | U-A | cWW | anti | anti | 10.957 | N3(imino)-N1[2.99]<br>O4(carbonyl)-N6(amino)[2.82] | Other |
| 4v6c | 1:DA.U2796 | 1:DA.A2800 | U-A | cWW | anti | anti | 10.68 | N3(imino)-N1[2.93]<br>O4(carbonyl)-N6(amino)[2.91] | Watson-Crick |
| 4v6d | 1:DA.U2796 | 1:DA.A2800 | U-A | cWW | anti | anti | 10.442 | N3(imino)-N1[2.90]<br>O4(carbonyl)-N6(amino)[2.92] | Watson-Crick |
| 4v6e | 1:DA.U2796 | 1:DA.A2800 | U-A | cWW | anti | anti | 10.217 | N3(imino)-N1[2.88]<br>O4(carbonyl)-N6(amino)[2.91] | Watson-Crick |
| 4v6f | 1:BA.U82 | 1:BA.A87 | U-A | cWW | anti | anti | 10.029 | N3(imino)-N1[2.78]<br>O4(carbonyl)-N6(amino)[2.98] | Watson-Crick |
| 4v6f | 1:CA.U82 | 1:CA.A87 | U-A | cWW | anti | anti | 10.486 | N3(imino)-N1[2.83]<br>O4(carbonyl)-N6(amino)[2.95] | Watson-Crick |
| 4v6g | 1:AA.U82 | 1:AA.A87 | U-A | cWW | anti | anti | 9.906 | N3(imino)-N1[2.78]<br>O4(carbonyl)-N6(amino)[2.97] | Other |
| 4v6g | 1:CA.U82 | 1:CA.A87 | U-A | cWW | anti | anti | 10.461 | N3(imino)-N1[2.82]<br>O4(carbonyl)-N6(amino)[2.95] | Watson-Crick |
| 4v7p | 1:DA.U82 | 1:DA.A87 | U-A | t.H | anti | anti | 10.179 | O2(carbonyl)-N6(amino)[3.68] | Other |
| 4v7p | 1:AA.U82 | 1:AA.A87 | U-A | t.H | anti | anti | 10.182 | O2(carbonyl)-N6(amino)[3.68] | Other |
| 4v7r | 1:C1.U1397 | 1:C1.A1401 | U-A | tWH | anti | anti | 10.663 | O2(carbonyl)-N6(amino)[3.37] | Other |
| 4v7r | 1:A1.U1397 | 1:A1.A1401 | U-A | tWH | anti | anti | 10.667 | O2(carbonyl)-N6(amino)[3.32] | Other |
| 4v7r | 1:D3.A109 | 1:D3.U113 | A-U | cWW | anti | anti | 10.677 | N6(amino)-O4(carbonyl)[2.97]<br>N1-N3(imino)[3.01] | Watson-Crick |
| 4v7r | 1:B3.A109 | 1:B3.U113 | A-U | cWW | anti | anti | 10.693 | N6(amino)-O4(carbonyl)[2.96]<br>N1-N3(imino)[3.01] | Watson-Crick |
| 4v7s | 1:DA.U2796 | 1:DA.A2800 | U-A | cWW | anti | anti | 10.679 | N3(imino)-N1[3.05]<br>O4(carbonyl)-N6(amino)[2.94] | Watson-Crick |
| 4v7t | 1:DA.U2796 | 1:DA.A2800 | U-A | cWW | anti | anti | 10.719 | N3(imino)-N1[3.01]<br>O4(carbonyl)-N6(amino)[3.01] | Watson-Crick |
| 4v7u | 1:DA.U2796 | 1:DA.A2800 | U-A | cWW | anti | anti | 11.3 | N3(imino)-N1[3.46]<br>O4(carbonyl)-N6(amino)[3.54] | Other |
| 4v7v | 1:DA.U2796 | 1:DA.A2800 | U-A | cWW | anti | anti | 10.786 | N3(imino)-N1[3.12]<br>O4(carbonyl)-N6(amino)[3.19] | Watson-Crick |
| 4v87 | 1:CA.U82 | 1:CA.A87 | U-A | cWW | anti | anti | 9.321 | N3(imino)-N1[2.64] | Other |

| | | | | | | | | | |
|---|---|---|---|---|---|---|---|---|---|
| | | | | | | | | O4(carbonyl)-N6(amino)[3.10] | |
| 4v87 | 1:BA.U82 | 1:BA.A87 | U-A | cWW | anti | anti | 10.05 | N3(imino)-N1[2.80]<br>O4(carbonyl)-N6(amino)[3.08] | Watson-Crick |
| 4v88 | 1:A5.U1567 | 1:A5.A1571 | U-A | cWW | anti | anti | 9.493 | N3(imino)-N1[3.22] | Other |
| 4v88 | 1:A6.U1397 | 1:A6.A1401 | U-A | tWH | anti | anti | 9.699 | O2(carbonyl)-N6(amino)[3.60] | Other |
| 4v88 | 1:A2.U1397 | 1:A2.A1401 | U-A | t.H | anti | anti | 10.092 | O2(carbonyl)-N6(amino)[3.60] | Other |
| 4v8b | 1:CA.U82 | 1:CA.A87 | U-A | cWW | anti | anti | 9.305 | N3(imino)-N1[2.69]<br>O4(carbonyl)-N6(amino)[3.17] | Other |
| 4v8b | 1:AA.U82 | 1:AA.A87 | U-A | cWW | anti | anti | 10.213 | N3(imino)-N1[2.80]<br>O4(carbonyl)-N6(amino)[3.07] | Watson-Crick |
| 4v8c | 1:DA.U82 | 1:DA.A87 | U-A | cWW | anti | anti | 9.278 | N3(imino)-N1[2.60]<br>O4(carbonyl)-N6(amino)[3.09] | Other |
| 4v8c | 1:CA.U82 | 1:CA.A87 | U-A | cWW | anti | anti | 10.136 | N3(imino)-N1[2.75]<br>O4(carbonyl)-N6(amino)[3.07] | Watson-Crick |
| 4v8d | 1:CA.U82 | 1:CA.A87 | U-A | cWW | anti | anti | 8.87 | O4(carbonyl)-N6(amino)[2.91] | Other |
| 4v8e | 1:DA.U82 | 1:DA.A87 | U-A | cWW | anti | anti | 9.293 | O2(carbonyl)*N1[2.15]<br>N3(imino)*N6(amino)[2.46]<br>N3(imino)-N1[2.83]<br>O4(carbonyl)-N6(amino)[2.98] | Other |
| 4v8f | 1:CA.U82 | 1:CA.A87 | U-A | cWW | anti | anti | 9.475 | O2(carbonyl)*N1[2.16]<br>N3(imino)*N6(amino)[2.43]<br>N3(imino)-N1[2.86]<br>O4(carbonyl)-N6(amino)[2.98] | Other |
| 4v8p | 1:F1.U1887 | 1:F1.A1891 | U-A | tSH | anti | anti | 9.151 | O2(carbonyl)-N6(amino)[3.07] | Other |
| 4v8p | 1:H1.U1887 | 1:H1.A1891 | U-A | tSH | anti | anti | 9.155 | O2(carbonyl)-N6(amino)[3.04] | Other |
| 4v8p | 1:D1.U1887 | 1:D1.A1891 | U-A | tSH | anti | anti | 9.173 | O2(carbonyl)-N6(amino)[3.10] | Other |
| 4v8p | 1:A1.U1887 | 1:A1.A1891 | U-A | tSH | anti | anti | 9.186 | O2(carbonyl)-N6(amino)[3.11] | Other |
| 4v9a | 1:CA.U82 | 1:CA.A87 | U-A | cWW | anti | anti | 8.681 | N3(imino)-N1[2.44]<br>O4(carbonyl)-N6(amino)[2.84] | Other |
| 4v9b | 1:CA.U82 | 1:CA.A87 | U-A | cWW | anti | anti | 8.969 | N3(imino)-N1[2.34]<br>O4(carbonyl)-N6(amino)[2.67] | Other |
| 4v9f | 1:0.U125 | 1:0.A129 | U-A | cWW | anti | anti | 10.584 | N3(imino)-N1[2.92]<br>O4(carbonyl)-N6(amino)[3.14] | Watson-Crick |
| 4v9j | 1:CA.U68^K | 1:CA.A68^O | U-A | tSH | anti | anti | 9.263 | O2'(hydroxyl)-N6(amino)[2.93]<br>O2(carbonyl)-N6(amino)[3.43] | Other |
| 4v9j | 1:AA.U68^K | 1:AA.A68^O | U-A | tSH | anti | anti | 9.503 | O2'(hydroxyl)-N6(amino)[3.08]<br>O2(carbonyl)-N6(amino)[3.54] | Other |
| 4v9k | 1:AA.U68^K | 1:AA.A68^O | U-A | c.W | anti | anti | 10.557 | O2(carbonyl)-N6(amino)[3.44] | Other |

| | | | | | | | | | |
|---|---|---|---|---|---|---|---|---|---|
| 4w29 | 1:CA.U68^K | 1:CA.A68^O | U-A | tSH | anti | anti | 9.186 | O2(carbonyl)-N6(amino)[3.47] | Other |
| 4w29 | 1:AA.U68^K | 1:AA.A68^O | U-A | t.H | anti | anti | 10.771 | O2(carbonyl)-N6(amino)[3.34] | Watson-Crick |
| 4wra | 1:1G.U82 | 1:1G.A87 | U-A | cWW | anti | anti | 9.869 | O2(carbonyl)*N1[2.63]<br>N3(imino)-N1[2.67]<br>O4(carbonyl)-N6(amino)[3.04] | Other |
| 4www | 1:YA.U2796 | 1:YA.A2800 | U-A | cWW | anti | anti | 10.658 | N3(imino)-N1[3.21]<br>O4(carbonyl)-N6(amino)[3.16] | Watson-Crick |
| 4x65 | 1:A.U81 | 1:A.A88 | U-A | t.H | anti | anti | 9.935 | O2(carbonyl)-N6(amino)[3.57] | Other |
| 4yy3 | 1:A.U81 | 1:A.A88 | U-A | t.H | anti | anti | 9.746 | O2(carbonyl)-N6(amino)[3.19] | Other |
| 5b2o | 1:B.A64 | 1:B.U68 | A-U | cWW | anti | anti | 10.29 | N6(amino)-O4(carbonyl)[3.12]<br>N1-N3(imino)[2.74] | Watson-Crick |
| 5b2o | 1:B.U82 | 1:B.A86 | U-A | cWW | anti | anti | 10.378 | N3(imino)-N1[2.78]<br>O4(carbonyl)-N6(amino)[3.08] | Watson-Crick |
| 5b2p | 1:B.U82 | 1:B.A86 | U-A | cWW | anti | anti | 10.286 | N3(imino)-N1[2.72]<br>O4(carbonyl)-N6(amino)[2.99] | Watson-Crick |
| 5b2p | 1:B.A64 | 1:B.U68 | A-U | cWW | anti | anti | 10.296 | N6(amino)-O4(carbonyl)[3.11]<br>N1-N3(imino)[2.75] | Watson-Crick |
| 5b2q | 1:B.A64 | 1:B.U68 | A-U | cWW | anti | anti | 10.289 | N6(amino)-O4(carbonyl)[3.10]<br>N1-N3(imino)[2.75] | Watson-Crick |
| 5b2q | 1:B.U82 | 1:B.A86 | U-A | cWW | anti | anti | 10.311 | N3(imino)-N1[2.73]<br>O4(carbonyl)-N6(amino)[3.00] | Watson-Crick |
| 5br8 | 1:A.U81 | 1:A.A88 | U-A | tSH | anti | anti | 9.746 | O2(carbonyl)-N6(amino)[3.94] | Other |
| 5ccb | 1:N.U33 | 1:N.A37 | U-A | cWW | anti | anti | 10.708 | N3(imino)-N1[2.93]<br>O4(carbonyl)-N6(amino)[2.99] | Watson-Crick |
| 5ccx | 1:N.U33 | 1:N.A37 | U-A | cWW | anti | anti | 10.589 | N3(imino)-N1[2.91]<br>O4(carbonyl)-N6(amino)[3.00] | Watson-Crick |
| 5cd1 | 1:N.U33 | 1:N.A37 | U-A | cWW | anti | anti | 10.715 | N3(imino)-N1[3.21]<br>O4(carbonyl)-N6(amino)[3.41] | Watson-Crick |
| 5d0b | 1:F.U33 | 1:F.A37 | U-A | cWW | anti | anti | 10.763 | N3(imino)-N1[3.11]<br>O4(carbonyl)-N6(amino)[3.20] | Watson-Crick |
| 5dat | 1:2.U1397 | 1:2.A1401 | U-A | tWH | anti | anti | 10.287 | O2(carbonyl)-N6(amino)[2.72] | Other |
| 5dc3 | 1:1.U1567 | 1:1.A1571 | U-A | cSW | anti | anti | 8.393 | O2(carbonyl)-N6(amino)[2.44] | Other |
| 5dc3 | 1:5.U1567 | 1:5.A1571 | U-A | cWW | anti | anti | 10.111 | N3(imino)-N1[3.51] | Other |
| 5dc3 | 1:2.U1397 | 1:2.A1401 | U-A | tWH | anti | anti | 10.228 | O2(carbonyl)-N6(amino)[2.71] | Other |
| 5dge | 1:5.U1567 | 1:5.A1571 | U-A | cWW | anti | -- | 10.349 | N3(imino)-N1[3.59] | Other |
| 5dgf | 1:2.U1397 | 1:2.A1401 | U-A | tWH | anti | anti | 10.165 | O2(carbonyl)-N6(amino)[2.44] | Other |
| 5dgv | 1:6.U1397 | 1:6.A1401 | U-A | tWH | anti | anti | 9.899 | O2(carbonyl)-N6(amino)[3.61] | Other |

| | | | | | | | | | |
|---|---|---|---|---|---|---|---|---|---|
| 5dgv | 1:2.U1397 | 1:2.A1401 | U-A | tWH | anti | anti | 10.451 | O2(carbonyl)-N6(amino)[2.75] | Other |
| 5dgv | 1:5.U1567 | 1:5.A1571 | U-A | cWW | anti | anti | 10.498 | N3(imino)-N1[3.95] | Other |
| 5fci | 1:5.U1567 | 1:5.A1571 | U-A | cWW | anti | anti | 9.739 | N3(imino)-N1[3.10] | Other |
| 5fcj | 1:5.U1567 | 1:5.A1571 | U-A | cWW | anti | anti | 9.693 | N3(imino)-N1[3.12] | Other |
| 5i4l | 1:5.U1567 | 1:5.A1571 | U-A | cWW | anti | anti | 9.84 | N3(imino)-N1[3.31] | Other |
| 5i4l | 1:2.U1397 | 1:2.A1401 | U-A | tWH | anti | anti | 9.965 | O2(carbonyl)-N6(amino)[3.31] | Other |
| 5lyb | 1:5.U1567 | 1:5.A1571 | U-A | cWW | anti | anti | 9.899 | N3(imino)-N1[3.47] | Other |
| 5lyb | 1:2.U1397 | 1:2.A1401 | U-A | t.H | anti | anti | 10.013 | O2(carbonyl)-N6(amino)[3.62] | Other |
| 5mei | 1:A.U1397 | 1:A.A1401 | U-A | tWH | anti | anti | 10.198 | O2(carbonyl)-N6(amino)[3.12] | Other |
| 5ndg | 1:6.U1397 | 1:6.A1401 | U-A | tWH | anti | anti | 9.981 | O2(carbonyl)-N6(amino)[3.18] | Other |
| 5ndg | 1:2.U1397 | 1:2.A1401 | U-A | tWH | anti | anti | 10.99 | O2(carbonyl)-N6(amino)[3.50] | Other |
| 5ndv | 1:2.U1397 | 1:2.A1401 | U-A | tWH | anti | anti | 9.453 | O2(carbonyl)-N6(amino)[2.59] | Other |
| 5ndv | 1:6.U1397 | 1:6.A1401 | U-A | tWH | anti | anti | 9.86 | O2(carbonyl)-N6(amino)[2.35] | Other |
| 5ndw | 1:6.U1397 | 1:6.A1401 | U-A | tWH | anti | anti | 10.19 | O2(carbonyl)-N6(amino)[3.17] | Other |
| 5ndw | 1:2.U1397 | 1:2.A1401 | U-A | t.H | anti | anti | 10.662 | O2(carbonyl)-N6(amino)[3.67] | Other |
| 5obm | 1:6.U1397 | 1:6.A1401 | U-A | tWH | anti | anti | 9.674 | O2(carbonyl)-N6(amino)[2.41] | Other |
| 5obm | 1:2.U1397 | 1:2.A1401 | U-A | tWH | anti | anti | 10.162 | O2(carbonyl)-N6(amino)[2.46] | Other |
| 5on6 | 1:A.U1397 | 1:A.A1401 | U-A | tWH | anti | anti | 10.203 | O2(carbonyl)-N6(amino)[3.35] | Other |
| 5tbw | 1:A.U1397 | 1:A.A1401 | U-A | tWH | anti | anti | 10.208 | O2(carbonyl)-N6(amino)[3.24] | Other |
| 5tga | 1:5.U1567 | 1:5.A1571 | U-A | cWW | anti | anti | 9.842 | N3(imino)-N1[3.35] | Other |
| 5tga | 1:2.U1397 | 1:2.A1401 | U-A | tWH | anti | anti | 10.036 | O2(carbonyl)-N6(amino)[3.02] | Other |
| 5tgm | 1:6.U1397 | 1:6.A1401 | U-A | tWH | anti | anti | 9.614 | O2(carbonyl)-N6(amino)[3.73] | Other |
| 5tgm | 1:2.U1380 | 1:2.A1384 | U-A | t.H | anti | anti | 9.894 | O2'(hydroxyl)-N6(amino)[3.43]<br>O2(carbonyl)-N6(amino)[3.57] | Other |
| 6bjx | 1:B.A235 | 1:B.U239 | A-U | cWW | anti | anti | 10.777 | N6(amino)-O4(carbonyl)[2.27]<br>N1-N3(imino)[2.52] | Watson-Crick |
| 6bjx | 1:A.A235 | 1:A.U239 | A-U | cWW | anti | anti | 10.814 | N6(amino)-O4(carbonyl)[2.23]<br>N1-N3(imino)[2.48] | Watson-Crick |
| 6cao | 1:A.U81 | 1:A.A88 | U-A | tWH | anti | anti | 10.633 | O2(carbonyl)-N6(amino)[3.30] | Other |
| 6cap | 1:A.U81 | 1:A.A88 | U-A | t.H | anti | anti | 10.518 | O2(carbonyl)-N6(amino)[3.89] | Other |

| | | | | | | | | | |
|---|---|---|---|---|---|---|---|---|---|
| 6cas | 1:A.U81 | 1:A.A88 | U-A | t.H | anti | anti | 10.297 | O2(carbonyl)-N6(amino)[3.59] | Other |
| 6d8l | 1:A.A235 | 1:A.U239 | A-U | cWW | anti | anti | 10.83 | N6(amino)-O4(carbonyl)[2.33]<br>N1-N3(imino)[2.64] | Watson-Crick |
| 6d8l | 1:B.A235 | 1:B.U239 | A-U | cWW | anti | anti | 10.932 | N6(amino)-O4(carbonyl)[2.37]<br>N1-N3(imino)[2.76] | Other |
| 6d8m | 1:B.A235 | 1:B.U239 | A-U | cWW | anti | anti | 10.905 | N6(amino)-O4(carbonyl)[2.83]<br>N1-N3(imino)[2.83] | Other |
| 6d8m | 1:A.A235 | 1:A.U239 | A-U | cWW | anti | anti | 11.069 | N6(amino)-O4(carbonyl)[2.75]<br>N1-N3(imino)[2.89] | Other |
| 6d8n | 1:A.A236 | 1:A.U240 | A-U | cWW | anti | anti | 10.778 | N6(amino)-O4(carbonyl)[2.39]<br>N1-N3(imino)[2.62] | Watson-Crick |
| 6d8n | 1:B.A236 | 1:B.U240 | A-U | cWW | anti | anti | 10.879 | N6(amino)-O4(carbonyl)[2.59]<br>N1-N3(imino)[2.79] | Other |
| 6d8o | 1:B.A235 | 1:B.U239 | A-U | cWW | anti | anti | 10.961 | N6(amino)-O4(carbonyl)[2.86]<br>N1-N3(imino)[2.98] | Other |
| 6d8o | 1:A.A235 | 1:A.U239 | A-U | cWW | anti | anti | 11.443 | N6(amino)-O4(carbonyl)[2.33]<br>N1-N3(imino)[3.02] | Other |
| 6fq3 | 1:B.U5 | 1:B.A9 | U-A | cWW | anti | -- | 10.395 | N3(imino)-N1[2.76]<br>O4(carbonyl)-N6(amino)[3.08] | Other |
| 6fql | 1:B.U5 | 1:B.A9 | U-A | cWW | anti | -- | 10.405 | N3(imino)-N1[2.86]<br>O4(carbonyl)-N6(amino)[3.20] | Other |
| 6gsl | 1:13.U82 | 1:13.A87 | U-A | cWW | anti | anti | 10.525 | N3(imino)-N1[2.83]<br>O4(carbonyl)-N6(amino)[2.82] | Watson-Crick |
| 6hhq | 1:sR.U1397 | 1:sR.A1401 | U-A | tSH | anti | anti | 9.727 | O2(carbonyl)-N6(amino)[3.77] | Other |
| 6ltp | 1:B.U12 | 1:B.A16 | U-A | cWW | anti | anti | 10.128 | N3(imino)-N1[2.65]<br>O4(carbonyl)-N6(amino)[2.98] | Watson-Crick |
| 6ltp | 1:H.U12 | 1:H.A16 | U-A | cWW | anti | anti | 10.305 | N3(imino)-N1[2.81]<br>O4(carbonyl)-N6(amino)[3.03] | Watson-Crick |
| 6ltr | 1:B.U12 | 1:B.A16 | U-A | cWW | anti | anti | 10.286 | N3(imino)-N1[2.74]<br>O4(carbonyl)-N6(amino)[2.94] | Watson-Crick |
| 6ltu | 1:B.U12 | 1:B.A16 | U-A | cWW | anti | anti | 10.294 | N3(imino)-N1[2.91]<br>O4(carbonyl)-N6(amino)[3.21] | Watson-Crick |
| 6lu0 | 1:B.U12 | 1:B.A16 | U-A | cWW | anti | anti | 10.351 | N3(imino)-N1[2.75]<br>O4(carbonyl)-N6(amino)[2.87] | Watson-Crick |
| 6n5n | 1:A.A15 | 1:A.U19 | A+U | cHW | syn | anti | 10.285 | N6(amino)-O4(carbonyl)[3.20] | Other (Hoogsteen-like) |
| 6qnr | 1:13.U82 | 1:13.A87 | U-A | cWW | anti | anti | 9.116 | N3(imino)-N1[3.35] | Other |
| 6tqa | 1:F.A9 | 1:F.U13 | A-U | cWW | anti | anti | 10.097 | N6(amino)-O4(carbonyl)[3.28]<br>N1-N3(imino)[2.97] | Watson-Crick |
| 6tqa | 1:G.A9 | 1:G.U13 | A-U | cWW | anti | anti | 10.178 | N6(amino)-O4(carbonyl)[3.02] | Watson-Crick |

| | | | | | | | | | |
|---|---|---|---|---|---|---|---|---|---|
| | | | | | | | | N1-N3(imino)[2.84] | |
| 6tqa | 1:E.A9 | 1:E.U13 | A-U | cWW | anti | anti | 10.195 | N6(amino)-O4(carbonyl)[3.14] N1-N3(imino)[2.90] | Watson-Crick |
| 6tqa | 1:H.A9 | 1:H.U13 | A-U | cWW | anti | anti | 10.25 | N6(amino)-O4(carbonyl)[3.24] N1-N3(imino)[2.85] | Watson-Crick |
| 6tqb | 1:B.A10 | 1:B.U14 | A-U | cWW | anti | anti | 10.088 | N6(amino)-O4(carbonyl)[3.14] N1-N3(imino)[2.88] | Watson-Crick |
| 7azs | 1:16SA.U725 | 1:16SA.A729 | U-A | cWW | anti | anti | 9.031 | N3(imino)-N1[3.30] | Other |
| 7dug | 1:A.U81 | 1:A.A88 | U-A | t.H | anti | anti | 10.21 | O2(carbonyl)-N6(amino)[3.63] | Other |
| 7duk | 1:A.U81 | 1:A.A88 | U-A | t.H | anti | anti | 10.343 | O2(carbonyl)-N6(amino)[3.51] | Other |
| 7dul | 1:A.U81 | 1:A.A88 | U-A | tWH | anti | anti | 10.42 | O2(carbonyl)-N6(amino)[3.59] | Other |
| 7osa | 1:18S.U1397 | 1:18S.A1401 | U-A | tWH | anti | anti | 10.574 | O2(carbonyl)-N6(amino)[3.07] | Other |
| 8c3a | 1:CM.U1383 | 1:CM.A1387 | U-A | tSH | anti | anti | 9.293 | O2'(hydroxyl)-N6(amino)[3.17] O2(carbonyl)-N6(amino)[3.60] | Other |
| 8c3a | 1:AS.U1859 | 1:AS.A1863 | U-A | tSH | anti | anti | 9.32 | O2(carbonyl)-N6(amino)[3.11] | Other |
| 8c3a | 1:1.U1859 | 1:1.A1863 | U-A | tSH | anti | anti | 9.359 | O2(carbonyl)-N6(amino)[3.00] | Other |
| 8c3a | 1:B.U1383 | 1:B.A1387 | U-A | tSH | anti | anti | 9.534 | O2(carbonyl)-N6(amino)[3.79] | Other |

**Supplementary References**


1. Shi, H., et al., Revealing A-T and G-C Hoogsteen base pairs in stressed protein-bound duplex DNA. *Nucleic Acids Res* **2021,** *49* (21), 12540-12555.
2. Adams, P. D., et al., PHENIX: a comprehensive Python-based system for macromolecular structure solution. *Acta Crystallogr D Biol Crystallogr* **2010,** *66* (Pt 2), 213-21.
3. Williams, C. J., et al., MolProbity: More and better reference data for improved all-atom structure validation. *Protein Sci* **2018,** *27* (1), 293-315.
4. Schanda, P., et al., SOFAST-HMQC experiments for recording two-dimensional heteronuclear correlation spectra of proteins within a few seconds. *J Biomol NMR* **2005,** *33* (4), 199-211.
5. Delaglio, F., et al., NMRPipe: a multidimensional spectral processing system based on UNIX pipes. *J Biomol NMR* **1995,** *6* (3), 277-93.
6. Lee, W., et al., NMRFAM-SPARKY: enhanced software for biomolecular NMR spectroscopy. *Bioinformatics* **2015,** *31* (8), 1325-7.
7. Nikolova, E. N., et al., Probing transient Hoogsteen hydrogen bonds in canonical duplex DNA using NMR relaxation dispersion and single-atom substitution. *J Am Chem Soc* **2012,** *134* (8), 3667-70.
8. Nikolova, E. N., et al., Transient Hoogsteen base pairs in canonical duplex DNA. *Nature* **2011,** *470* (7335), 498-502.
9. Hansen, A. L., et al., Extending the range of microsecond-to-millisecond chemical exchange detected in labeled and unlabeled nucleic acids by selective carbon R(1rho) NMR spectroscopy. *Journal of the American Chemical Society* **2009,** *131* (11), 3818-9.
10. Kimsey, I. J., et al., Visualizing transient Watson-Crick-like mispairs in DNA and RNA duplexes. *Nature* **2015,** *519* (7543), 315-20.
11. Rangadurai, A., et al., Characterizing micro-to-millisecond chemical exchange in nucleic acids using off-resonance R(1rho) relaxation dispersion. *Prog Nucl Magn Reson Spectrosc* **2019,** *112-113*, 55-102.
12. McConnell, H. M., Reaction Rates by Nuclear Magnetic Resonance. *The Journal of Chemical Physics* **1958,** *28* (3), 430-431.
13. Bothe, J. R., et al., Evaluating the uncertainty in exchange parameters determined from off-resonance R1rho relaxation dispersion for systems in fast exchange. *J Magn Reson* **2014,** *244*, 18-29.
14. Abramson, J., et al., Accurate structure prediction of biomolecular interactions with AlphaFold 3. *Nature* **2024,** *630* (8016), 493-500.
15. Berman, H., et al., Announcing the worldwide Protein Data Bank. *Nat Struct Biol* **2003,** *10* (12), 980.
16. Lu, X. J.; Olson, W. K., 3DNA: a software package for the analysis, rebuilding and visualization of three-dimensional nucleic acid structures. *Nucleic Acids Res* **2003,** *31* (17), 5108-21.
17. Leontis, N. B.; Westhof, E., Geometric nomenclature and classification of RNA base pairs. *RNA* **2001,** *7* (4), 499-512.
18. Johnson, M., et al., NCBI BLAST: a better web interface. *Nucleic Acids Res* **2008,** *36* (Web Server issue), W5-9.
19. Eaton, K. A., et al., Role of Helicobacter pylori cag region genes in colonization and gastritis in two animal models. *Infect Immun* **2001,** *69* (5), 2902-8.
